# Mammalian Amoeboid Swimming is propelled by molecular and not protrusion-based paddling in Lymphocytes

**DOI:** 10.1101/509182

**Authors:** Laurene Aoun, Paulin Nègre, Alexander Farutin, Nicolas Garcia-Seyda, Mohd Suhail Rizvi, Rémi Galland, Alphée Michelot, Xuan Luo, Martine Biarnes-Pelicot, C. Hivroz, Salima Rafai, Jean-Baptiste Sibarita, Marie-Pierre Valignat, Chaouqi Misbah, Olivier Theodoly

## Abstract

Mammalian cells developed two main migration modes. The slow mesenchymatous mode, like fibroblasts crawling, relies on maturation of adhesion complexes and actin fiber traction, while the fast amoeboid mode, observed exclusively for leukocytes and cancer cells, is characterized by weak adhesion, highly dynamic cell shapes, and ubiquitous motility on 2D and in 3D solid matrix. In both cases, interactions with the substrate by adhesion or friction are widely accepted as a prerequisite for mammalian cell motility, which precludes swimming. We show here experimentally and computationally that leukocytes do swim, and that propulsion is not fueled by waves of cell deformation but by a rearward and inhomogeneous treadmilling of the cell envelope. We model the propulsion as a molecular paddling by transmembrane proteins linked to and advected by the actin cortex, whereas freely diffusing transmembrane proteins hinder swimming. This mechanism explains that swimming is five times slower than the cortex retrograde flow. Resultantly the ubiquitous ability of mammalian amoeboid cells to migrate in various environments can be explained for lymphocytes by a single machinery of envelope treadmilling.

---

Individual living cells evolved different strategies to migrate and explore their environment. Bacteria, microalgae or mammalian gametes can swim in suspension in a fluid, under the propulsion of a flagellum^1^ or of shape deformations^2^, whereas somatic mammalian cells crawl with adhesion on a solid tissue, via a continuous sequence of forward pushing of the cell front, strengthening of adhesions at the leading edge, and pulling of the cell rear^3,4^. In vivo, mammalian cells crawl either on 2D substrates, like leukocytes on inner blood vessels or epithelial surfaces, or in 3D environments within tissues. The critical role of adhesion for crawling motility was recently revised in the case of amoeboid mammalian cells, i.e. white blood cells and cancer cells. Amoeboid cells differ from mesenchymatous cells (e.g. fibroblasts) by a significantly higher velocity (typically 5-20 μm.min^−1^ vs 0. 1-1 μm.min^−1^) and highly dynamic shape deformations. Whereas amoeboid cells crawl on adhering substrates like mesenchymatous cells do, they also remain highly motile in the absence of adhesion, provided that they are confined by a 3D environment^5–7^. This motility in non-adherent conditions was explained by a chimneying^6^ mode where cell-substrate interactions are mediated by friction instead of adhesion^8–10^. Altogether, there is common agreement that amoeboid motility of mammalian cells is strictly dependent on adhesion on 2D substrates and on adhesion/friction in 3D media, while non-adherent 2D migration and swimming are precluded^5,11^.

In contradiction with the paradigm of adhesive or frictional crawling, Barry and Bretscher^12^ reported in 2010 that human neutrophils do swim. They suggested that propulsion may result from membrane treadmilling (rearward movement of the cell surface) or shape-deformation (protrusions and contractions along the cell body) but provided no experimental or theoretical evidence supporting either of these hypotheses. Most investigations were later performed on swimming of a non-mammalian eukaryotic cell, the amoeba *Dyctyostellium discoideum*. Some studies have defended a deformation-based propulsion^13,14^, whereas another one discarded both treadmilling and shape deformation^15^. For tumoral cells, one theoretical model of blebbing mentioned the possibility of migration in suspension by shape changes^16^, whereas other modeling efforts validated a swimming mechanism based on shape deformation for the case of cyanobacteria^17^ and microalgae^2^. A recent study on mesenchymatous macrophages cell line RAW 264.7 reported an amoeboid swimming mode artificially triggered by optogentic activation of actomyosin contractility in cell rear^18^. Propulsion convincingly involved membrane treadmilling, whereas contribution deformations was not assessed. Altogether, swimming of cells without flagellum remains mostly explained by shape deformation mechanisms, and besides, swimming of mammalian cells without flagellum remains widely discarded^5,6,11,19–23^.

Here, we demonstrate the existence of mammalian amoeboid swimming on human T lymphocytes and decipher its functioning experimentally and theoretically at the cellular and molecular scales. T lymphocytes are known to crawl on 2D adhering substrates^24–27^ and in 3D matrices via adhesion/friction^10,28,29^ at typical velocities of 20 μm.min^−1^. We observed swimming both in bulk solution and in the vicinity of anti-adhesive substrates and quantified an average speed of 5 μm.min^−1^. Experimental and theoretical evidences show that swimming is propelled by a molecular paddling of the transmembrane proteins linked to actin, which are axisymmetrically recycled between cell front and back by retrograde treadmilling at the plasma membrane and anterograde vesicular transport inside cells. This molecular description explains also quantitatively why swimming is significantly slower than the retrograde flow of the cell cortex and cell adhesive crawling.

## RESULTS

### Leukocytes swim without adhesion or friction with a solid substrate

Upon recruitment from the blood stream toward inflammation zones, leukocytes arrest and crawl on the inner surface of blood vessels (2D migration). Crawling was here reproduced in vitro with human primary effector T lymphocytes on glass substrates coated with ICAM-1 molecules, a specific ligand for the integrin adhesion molecules LFA-1 (αLβ2). Effector T lymphocytes were already polarized in suspension, with a front pole forming protrusions under the influence of actin polymerization and a rear pole undergoing contraction cycles enforced by acto-myosin contractility. When introduced into a chamber coated with ICAM-1, lymphocytes sedimented, adhered to, and migrated on the substrates. They crawled with a random walk pattern (Figure 1-A and Suppl. Mat. Movie 1) of curvilinear velocity 14.7 ± SD 7.5 μm.min^−1^, with a wide front pole (lamellipod), the nucleus positioned in the cell central zone, and a narrow tail (uropod). Interferometric imaging, in which a dark contrast of adhesive zones corresponds to a distance smaller than 50 nm^30^, attested for a molecular contact between cell and substrate (Figure 1-A-iii and Suppl. Mat. Movie 1). To challenge the idea that adhesion is necessary for amoeboid migration on a 2D substrate, we replaced the ICAM-1 surface treatment by an anti-adhesive coating of Pluronic^©^ F127. In the absence of adhesion cells were highly sensitive to residual flow drifts, so the determination of intrinsic self-propulsion required experimental caution to approach “zero flow” conditions. Using narrow microfluidic channels and pressure controllers (see material and methods) to reduce lateral drift, we observed cell sedimentation on the substrate and a random walk migration (Figure 1-B and Suppl. Mat. Movie 1) with an apparent average curvilinear speed of 5.5 ± SD 2.2 μm.min^−1^. The shape of the swimming cells (Figure 1-B-ii and Suppl. Mat Movie 1) looked similar to the ones of crawling cells with a somewhat less wide lamellipod. However, interferometric imaging attested that swimming cells were non-adherent (Figure 1-B-iii and Suppl. Mat. Movie 1), as the brightness of the contact zone corresponded to a liquid film separating the cell membrane from the substrate by more than 100 nm. Swimming in the vicinity of a substrate was further imaged in 3D by spinning-disk microscopy (Suppl. Mat. Movie 2) and, although strong phototoxicity hampered long-term 3D imaging, swimming of polarized cells with highly dynamic 3D shape deformation was evident on tens of micrometers. Hence, in contrast to most literature reports, lymphocytes do migrate on a 2D surface in the absence of adhesion.

**Figure 1:**
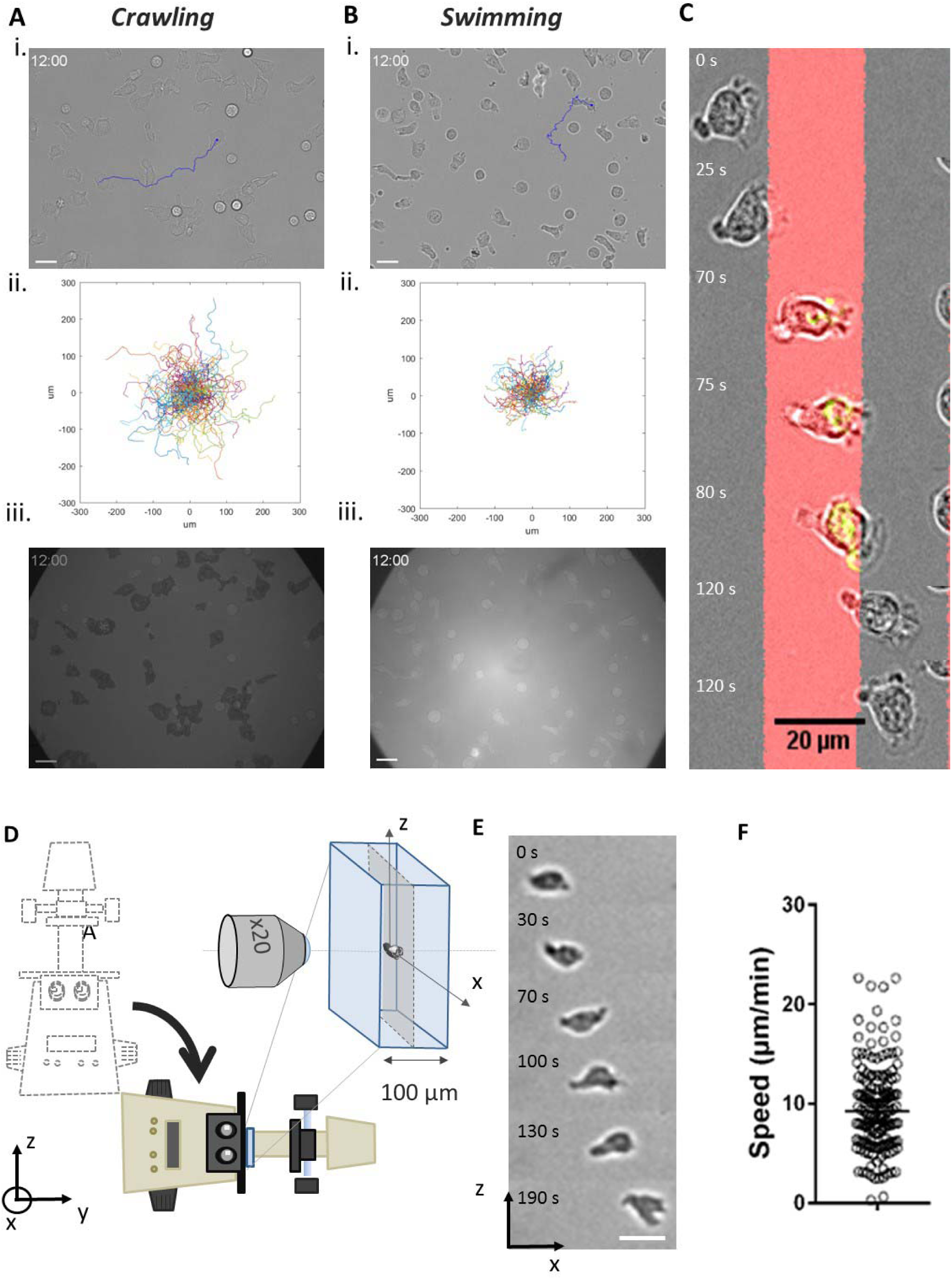
Crawling to Swimming in 2D and 3D displays a single motility mode. Primary human effector T cells **(A)** crawling on adhesive ICAM-1-treated substrate, and **(B)** swimming on an anti-adhesive Pluronic F127-treated substrate; **i.** 63x Bright field images. Blue line correspond to the track of one particular cell (time lag 10 s, duration 12 mn). Scale bar is 20 μm. **ii.** Representative tracks of motile cells in a single experiment (time lag = 20 s, duration 16 mn, n > 100 cells), **iii.** 63x Reflective Interference Contrast Microscopy (RICM) images. Cell contact zone is dark for cells crawling on ICAM treated surfaces, revealing an adhesion phenotype (left), and white for cells on non-adhesive surface attesting the absence of adhesion to the surface. Scale bar is 20 μm. **(C) No apparent transition between crawling and swimming.** Image sequence of cells migrating over adjacent stripes of adhesive ICAM-1 and anti-adhesive Polyetheleneglycol prepared by LIMAP^54^ with a width of 20 μm. The sequence is a merge of fluorescent images (ICAM-1, red), bright field images (greyscale) and reflection interference contrast microscopy images (adhesion zone, green). Scale bar 20 μm. **(D-E) Swimming in free suspension.** (D) Schematic of the set-up used for suspension swimming analysis with a microscope tilted by 90° and the flow channel oriented vertically allowing sideway observation. (E) Sequence of images of a cell swimming in the center of the channel along axis x. Scale bar 20 μm. **(F)** Velocities of swimming cells for a distance to wall larger than 40 μm. Nexperiments = 10, Ncells = 15, Nsteps > 200.

### Transition between crawling and swimming is fast

Swimming appears slower than crawling, however it is not clear if this difference results from the existence of two distinct machineries or only from a different coupling between the environment a conserved propelling machinery. To shed light on a potential switch between distinct migration modes, we presented cells to substrates patterned with alternated stripes of adhesive and anti-adhesive coatings with a periodicity of 20 μm (Figure 1-C and Suppl. Mat. Movie 3). Surprisingly, cells escaped frequently the adhesive zones (red stripes), something not observed in similar experiments with mesenchymatous cells^31^. Furthermore, although interference microscopy attested that cells travelled across the adhesive zone with adhesion (green signal) and on anti-adhesive zones without attachment (no green signal), there was no evident lag time or change in cell morphology dynamics upon transition from crawling to swimming. These observations support either that switching between crawling/swimming modes is fast, or that crawling and swimming share a common machinery, discarding the existence of a mode switching.

### Leukocytes swim in free suspension

Although leukocytes migrated in the vicinity of a surface without adhesion, cell-substrate distance remained in the nanometric range. Hence, swimming close to a substrate could rely on hydrodynamic coupling between cell and substrate. We therefore performed experiments with cells in bulk suspension to avoid any hydrodynamic interference with solid walls. The experimental challenge consisted in cancelling all artefactual passive cell movements that may superimpose to active self-propulsion. Passive cell displacement can arise from cell sedimentation as well as from flow drifts due to temperature gradients, pressure imbalance between channel outlets or gravimetric imbalance due to cell dispersion inhomogeneity. Experiments were performed with a 90° tilted microscope, a vertical microfluidic channel, Ficoll supplemented medium to match the average density of cells, a high precision pressure controller and highly resistant tubing connections to slow down pressure-driven flows (Figure 1-D and Suppl. Mat. for details). In this configuration, the vertical axis corresponded to the direction of both pressure-driven flow across the microchannel and gravity-induced sedimentation flow. Hence, cell velocity along the vertical axis was not quantitatively exploitable, because flow drifts and sedimentation effects, although lowered, remained in the range of a few μm.min^−1^ and were not negligible as compared to swimming velocities (Figure 1-E and suppl. Mat. Movie 4). Swimming prowess was nevertheless measurable on the two other axes. Figure 1-F shows that cells in bulk suspension swam with an apparent average curvilinear speed around 9.5 ± SD 4.2 μm.min^−1^ (images taken every 30s), which confirmed the intrinsic capability of lymphocytes to swim. In what follows, systematic measurements were however performed with cells close to a non-adherent substrate because theoretical calculations agreed that the vicinity of a single wall has a negligible effect on swimming velocity (Figure S 5).

### Diffusive versus active motion in swimming conditions

Cells velocity estimated by averaging the displacements of cell mass centres over intervals of 30 s yielded a significant difference between swimming and crawling (**Figure 2**-A). However, the cell population obtained from the in vitro activation of lymphocytes comprises two fractions of cells, one of round and inactive cells, and the other of polarized and active cells. In crawling experiments, round cells, generally non-adherent, were washed away by residual flows and therefore not present in the recorded data. By contrast, in swimming experiments, both active and inactive cells were taken into account because residual flows were cancelled, so that the average raw velocity was biased towards lower values for swimming. **Figure 2**-B presents the histogram of raw velocities for each individual cells of a population of live lymphocytes and of the same population fixed by paraformaldehyde where all cells have a frozen shape. The histogram of live cells presents two populations. One population has a low velocity close to the one of fixed cells, allowing one to link this population with inactive cells. The high velocity population, corresponding to swimming cells, is characterized by a fraction of 80% and an average raw curvilinear speed of 5.9 ± 4.2 μm.min^−1^. However, this determination of swimming speed includes also from diffusion effects. Indeed, fixed cells without swimming activity have an average raw curvilinear speed of 2.8 ± 0.3 μm.min^−1^. We therefore performed a detailed analysis of the cell trajectories to extract diffusive contribution to motion. First, we investigated the mean square displacement averaged over all cells in the population as a function of time. Second, we fitted the mean square displacements as a function of time interval by a random-walk law, which combines 2D Brownian-like diffusion with persistent motion, and we analysed the distribution of velocities *v*_*s*_ and diffusion coefficients *D*_*t*_ obtained by the fitting procedure:

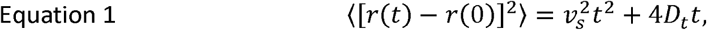

**Figure 2:**
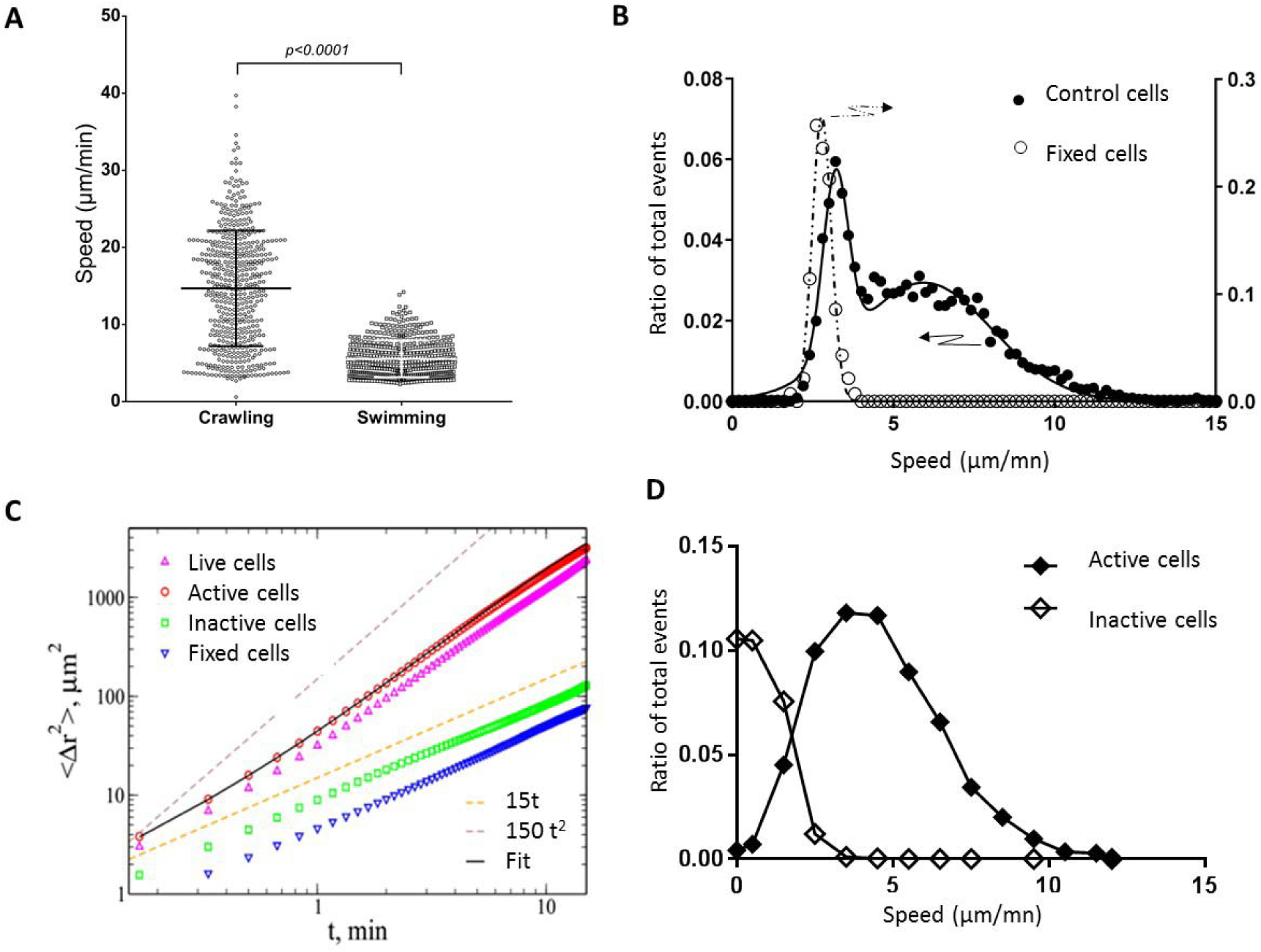
Active swimming propels cells at 5 μm.s^−1^. **(A)** Raw curvilinear speed of cells crawling on adherent ICAM-l-treated substrate and swimming on anti-adherent Pluronic F127-treated substrate, estimated by averaging the displacements of cell mass centres over intervals of 30 s. N = 500 cells, p value oft test. **(B)** Histogram of raw curvilinear speed of control swimming cells (filed dots) and fixed cells (hollow dots). Data are fitted with a single Gaussian for fixed cells (dotted line) and a double Gaussian for control cells (dark line). Live cells are composed of one population of diffusing cells and one of swimming cells with an average speed of 5.9 μm.min^−1^. **(C)** Mean square displacement < [*r*(*t*) *– r*(*0*)]^2^ > as a function of time for all cells and all steps combined in the case of live cells (upward pointed triangles) and fixed cells (downward pointed triangles). Fixed cells have purely diffusive behavior corresponding to *D*_*t*_ *=* 2.34 μm^2^.min^−1^. Circles and squares show the mean square displacement for active and inactive cells, respectively. Black line is a fit of active cells using Equation 9 (Suppl. Mat.) with *v*_*s*_ = 4.3 μm.^−1^, *D*_*r*_ = 0.19 min^−1^, and *D*_*t*_ = 7.28 μm^2^ min^−1^. **(D)** Histogram speed measured individually using Equation 1 as in (C) for active cells (filled diamonds) and inactive cells (hollow diamonds). Root mean square velocity of active cells is equal to 4.9 μm.min^−1^. Lines are guides for the eye.

We then separated all cells in the population into two groups. We considered active the cells that travelled at least a distance of 25 μm (about two cell diameters) during the acquisition time of 13 min. The rest of the cells were referred to as inactive. The results of the analysis are shown in **Figure 2-C and D**. **Figure** 2-C shows the mean square displacement as a function of time for all cells combined, and for inactive and active cells separately. We also present the results with fixed cells, which are only affected by thermal diffusion. The guides suggest that both inactive and fixed cells had a diffusive behaviour, with an average diffusion coefficients of respectively 2.3 μm^2^.min^−1^ and 1.1 μm^2^.min^−1^, respectively. On the contrary, active cells showed a superdiffusive behaviour, which could not be fitted by Equation 1. A satisfactory fit (black curve in Figure 2-C) was obtained when we extended the model by adding rotational diffusion, which accounted for gradual changes in the swimming direction of the cells (see Suppl. Mat.). The fitting procedure gave here *v*_*s*_ = 4.3 μm.min^−1^, *D*_*r*_ = 0.19 min^−1^ and *D*_*t*_ = 7.28 μm^2^.min^−1^. To obtain the distribution of velocities of active cells, we simplified the analysis and considered only displacements for time intervals of 2 min, which is 3 times smaller than <1/D_r_, to remove the influence of the rotational diffusion. As can be observed in **Figure** 2-D, most of the active cells had a velocity around 3 to 5 μm.min^−1^. Root mean square velocity extracted from individual fits of active cells ends up to be equal to 4.9 μm.min^−1^ in cell medium. In order to determine the influence of viscosity on the swimming speed, similar experiments and analysis were also performed in medium supplemented with dextran of molecular weight 2,000 kDa. Swimming speed was found unchanged when viscosity was increased 100 times (Suppl. Mat. Table 1 and Figure S 1). Altogether, the average swimming velocity is largely independent of the viscosity of the external medium and equal to around 5 μm. min^−1^, which is significantly smaller than the crawling speed of 15 μm. min^−1^, and.

### Actin mediates swimming propulsion by polymerization and to a lesser extent contractility

Actin cytoskeleton is widely accepted as the molecular engine that propels cell crawling. To get more insight into its role on cell swimming, we used several actin inhibitors. Effector T cells are characterized by a strongly polarized state with actomyosin contractility mainly in the cell rear (uropod), and actin polymerization mainly in the cell front (lamellipod). Blebbistatin, a potent inhibitor of actomyosin contractility, strongly affected cells morphology (Figure 3-A and Suppl. Mat. Movie 5), as cells displayed a roughly round cell body with no distinct uropod and no contractile activity. Active cells had nevertheless a small-size lamellipod, which attested that they conserved a partial, albeit stable front-rear polarization. Interestingly, active cells were still swimming, and always in the direction of the lamellipod. The velocity and fraction of swimming cells were decreased around a factor of two as compared to control cells (Figure 3-B and Table 1). These results prove that frontal polymerization alone can propel swimming, while rear contractility is not necessary although it somehow participates to propulsion efficiency. We then perturbed actin polymerization in the cell front with Latrunculin A (Figure 3-A Suppl. Mat. Movie 5). The dose was specially titrated to inhibit the lamellipod at the cell front while preserving contractility in the cell rear. Latrunculin-treated cells were deprived of lamellipod and conserved a uropod, which is the opposite situation to blebbistatin-treated cells. The fraction of swimming cells and the mean velocities were significantly lower than for blebbistatin-treated cells (Figure 3-B and Table 1), which supports further that the lamellipod plays a preponderant role in swimming propulsion. We then treated cells with CK666, an inhibitor of the protein Arp2/3 that mediates branching of the actin network in lamellipods (Figure 3-A Suppl. Mat. Movie 5). While the front of migrating leukocytes usually displays lamellar shaped protrusions^32^, CK666-treated cells formed filopodia and blebs in the cell front. The effect of CK666 on swimming speed was found intermediate between blebbistatin and Latrunculin cases. Altogether, swimming was more efficient with a perturbed lamellipod (CK666) than without lamellipod (Latrunculin), which is self-understanding since lamellipod was found important for propulsion. Moreover, swimming was more efficient for totally inhibited uropod (CK666) than for partially inhibited lamellipod (blebbistatin), which confirmed that swimming is mediated in a larger extent by lamellipod rather than uropod. Finally, swimming was fully abrogated with a combination of blebbistatin and Latrunculin (Figure 3-A,B), from which we conclude that the actin network is the only engine of lymphocyte swimming.

**Figure 3:**
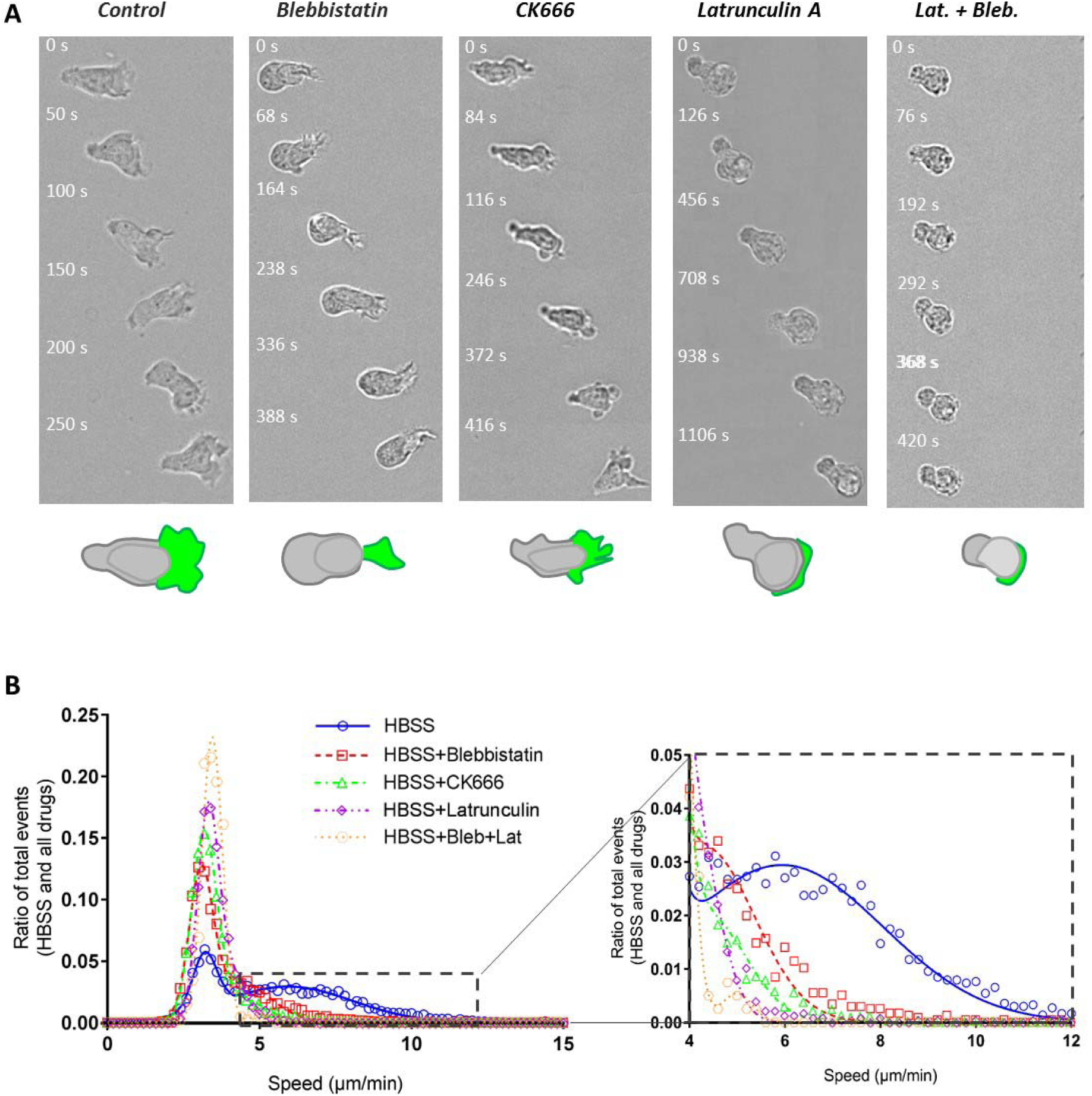
Both actin polymerization and contractility propel swimming. **(A)** Bright field image sequences showing the shape and dynamics of cells swimming on an antiadhesive substrate versus addition of actomyosin inhibitors. From left to right: wild type cells, cells treated with 50 μM blebbistatin, 100 μM CK666, 0.05 μM Latrunculin and combined Latrunculin and Blebbistatin. Cartoons at the bottom reproduce the cell in the first image to illustrate in each case the shape of the cell body (rear and nucleus) in grey and of the cell front or lamellipodium in green. Blebbistatin-treated cells have a roundish cell body without travelling protrusion, and a reduced but active lamellipod; CK666-treated cells have a perturbed lamellipod forming blebs and spikes; Latrunculin-treated cells have almost no lamellipod. Cells treated with Blebistatine and Latruculin have a roundish non contractile cell body and no lamellipod (scale bars: 50 μm top, 10 μm bottom). **(C)** Histogram of raw curvilinear velocities for swimming cells in response to above mentioned actin inhibitors treatments. Insert presents a zoom of the histogram corresponding to the active cells Gaussian. Ncells = 4342 (HBSS), 2353 (Blebbistatin), 5582 (CK666), 2255 (Latrunculin), 403 (Blebbistatine+Latrunculin); Nexperiments > 5.

**Table 1:** Swimming velocity versus the effects of cell fixation, medium viscosity change and actin inhibitors addition. The table reports the raw swimming velocity estimated by averaging the instant curvilinear speeds between two positions of cell mass center separated by 30s for each cell. Average raw velocities of active and passive cells as well as fraction of active cells are estimated from by a fit of velocities histograms by a double Gaussian (Figure 2-B). Velocity errors correspond to standard deviation. Cell number correspond to the number of cells considered for each experimental condition.

|  | Active cells |  | Passive cells |  |
| --- | --- | --- | --- | --- |
| Condition | ($\mu\text{m}.\text{min}^{-1}$) | Fraction | ($\mu\text{m}.\text{min}^{-1}$) | Number of cells |
| HBSS-CTRL | $5.9 \pm 2.1$ | 78% | $3.2 \pm 0.4$ | 4342 |
| Blebbistatin | $4.3 \pm 1.1$ | 45% | $3.1 \pm 0.4$ | 2353 |
| CK666 | $4.4 \pm 0.8$ | 21% | $3.2 \pm 0.4$ | 5582 |
| Latrunculin | $4.2 \pm 0.5$ | 20% | $3.3 \pm 0.4$ | 2255 |
| Bleb. + Lat. | $5.0 \pm 0.3$ | 2% | $3.4 \pm 0.3$ | 403 |
| HBSS-PFA | - | - | $2.8 \pm 0.3$ | 139 |
| Viscosity 10x | $4.4 \pm 1.6$ | NA | - | 1262 |
| Viscosity 100x | $6.0 \pm 2$ | NA | - | 64 |
| Dextrose | $5.9 \pm 1.8$ | 80% | $3.6 \pm 0.6$ | 1449 |
| Visco 10x-PFA | - | - | $0.27 \pm 0.09$ | 30 |
| ICAM-1 | $14.6 \pm 7.5$ | - | - | 503 |

### Paddling by rearward travelling of membrane protrusions does not propel lymphocyte swimming

Swimming propulsion arises from the interactions of the cell external envelope with the surrounding fluid, therefore the dynamic properties of the cell external envelope is the key of the swimming mechanism. Like all amoeboid cells, lymphocytes display highly dynamic shape deformations. These normal movements of cell envelope are therefore a good candidate for propulsion drive. Since spinning disk imaging was limited by the strong photosensitivity of primary human T lymphocytes, we resorted to light-sheet soSPIM microscopy and transfected cells with RFP-Lifeact to visualize precisely the 3D dynamics of cell cytoskeleton (Figure 4-A and Suppl. Mat. Movie 6). 3D imaging revealed waves of lamellar protrusions that formed in the cell front^32^ with different orientations, traveled backwards and vanished when reaching the cell central and rear zones. Similar propagating waves of cell envelope were also visible in transmission microscopy (**Figure 4**-B), together with constriction rings (**Figure 4**-C)^21^. Constriction rings formed around the cells central zone, and the nucleus was intermittently pushed forward through the rings, provoking sudden and important reorganization of the contour of cells (Suppl. Mat. **Movie** 7). Hence, the dynamics of lymphocyte morphology is qualitatively reminiscent of the shape-deformation cycles analyzed in theoretical modeling of amoeboid swimming^2^. However, Figure 4-D shows that blebbistatin-treated cells keep quasi-static cell body shapes, which is very different to control cells, but nevertheless swim. This observation strongly suggests that shape deformations is not essential in leukocyte swimming. To assess precisely its contribution, we developed a direct numerical simulation of normal active forces applied to the cell membrane (with force-free and torque-free conditions) that generated “blebbing waves” backwards along the cell body (Figure 4-E and suppl. Mat Movie 8). Simulations showed that deformation cycles yield swimming propulsion with a generated swimming motion about 1000 times slower than the blebbing wave. In contrast, the waves speed were found experimentally around 10 μm.min^−1^, which is in the same magnitude as the swimming speed (Figure 4). We conclude that amoeboid shape deformation, i.e. normal movements of cell envelope, cannot explain the amoeboid swimming of lymphocytes.

**Figure 4:**
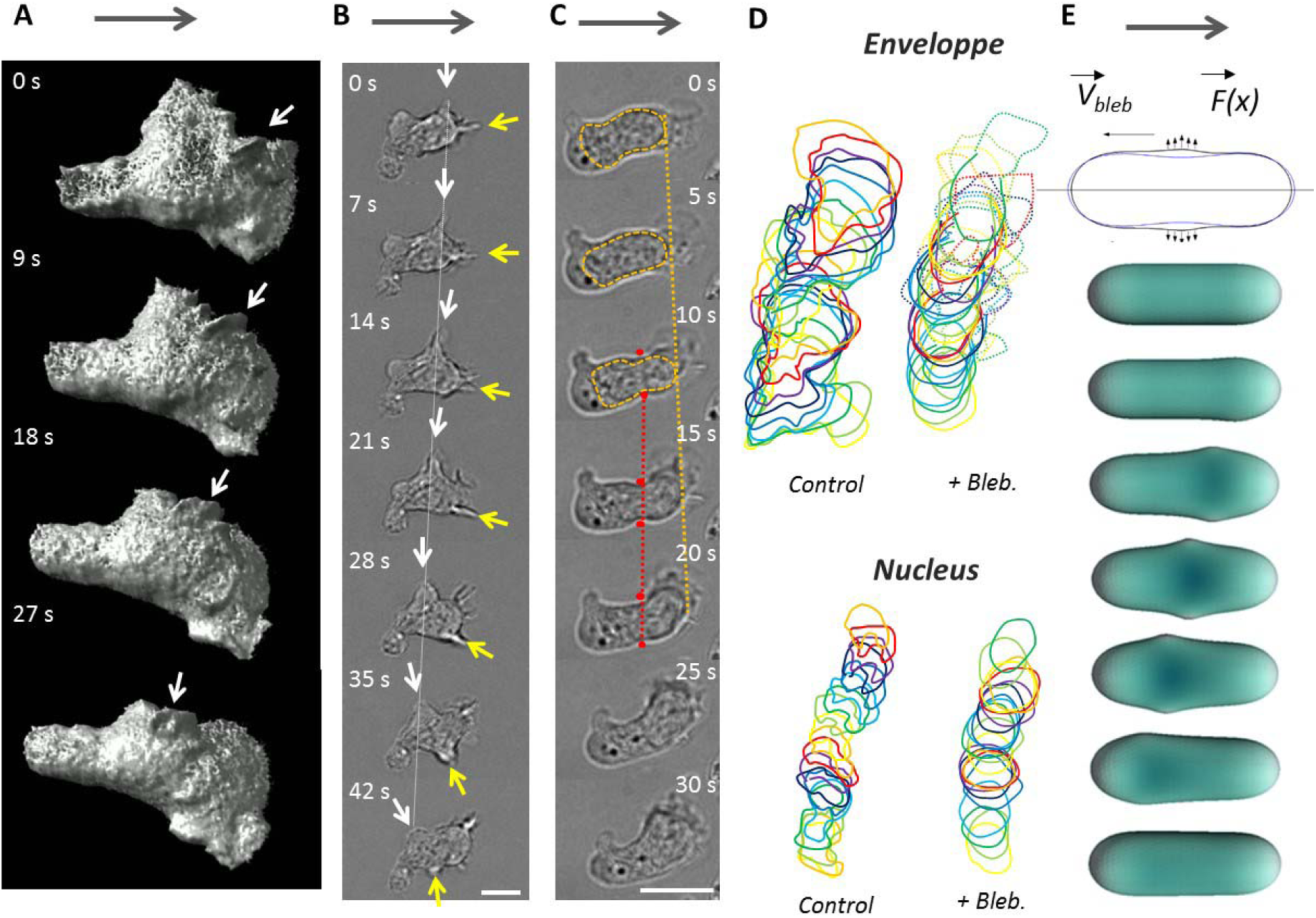
Protrusion paddling does not propel swimming. **(A-C)** Images sequence of swimming cell with micron scale protrusions travelling along cell body. **(A)** SoSPIM images of a cell transfected with RFP-Lifeact reveals the shape and motion of waves of actin protrusion in three dimensions. White arrow points to one particular protrusion. **(B,C)** Bright field images of a swimming cell showing dynamics of protrusions and constrictions along cell body. (B) Protrusions (white and yellow arrows) travel backwards in the frame of the cell and of the lab (white dashed line). (C) Constriction ring (between the two red dots) are hardly mobile in the frame of the lab (red dashed line), whereas the nucleus squeezing through the ring moves forward in the frame of the cell and of the laboratory (orange dashed line). Grey arrows indicate swimming direction. Scale bar is 10 μm. **(D)** Representative sequences of the contours of a cell (top) and its nucleus (bottom) for control cells (left) and blebbistatin-treated cells (right). For Blebbistatin, the envelope contour is a full line for cell rear body and dotted line for the lamellipod. The body and nucleus display strong deformation in control cells, and a quasi-static shapes with blebbistatin. Time lag is 10s for control and 15 s with blebbistatin. **(E)** Schematic illustrating the model of cell swimming by protrusive blebs. (Top) Blue and black contours are the initial and deformed configurations of the cell in the model. (Bottom) Sequence of cell shapes obtained by the numerical simulation. Simulation yield that a cell propelled by shape waves that is 1000 times slower than the protrusion wave (details in Suppl. Mat.).

### Membrane rearward treadmilling correlates with swimming speed

Normal motion of cell envelope does not yield sufficient propulsion, but the membrane of amoeboid cells displays also tangential movement, triggered by the retrograde flow of the inner actin cortex. To probe the motion of cells external envelope, we tracked beads coated by ICAM-1 molecules and attached to cells via the transmembrane integrins LFA-1 (Figure 5-A and Movie 9). Beads displayed a backward motion in the reference frame of the cell with an average velocity of 12 ± 3 urn.min^−1^, which is significantly larger than cell swimming velocity. The same experimental approach was applied in presence of the actin inhibitors blebbistatin, CK666 and latrunculin A (Figure 5-A and Movie 9). The retrograde motion of beads attached to cells membrane decreased in the presence of all inhibitors. Furthermore, the decrease of beads velocity with blebbistatin, CK666 and latrunculin A correlated with the respective decrease in swimming speed (Figure 5-B). These data strongly suggest a link between tangential envelope motion and swimming propulsion.

**Figure 5:**
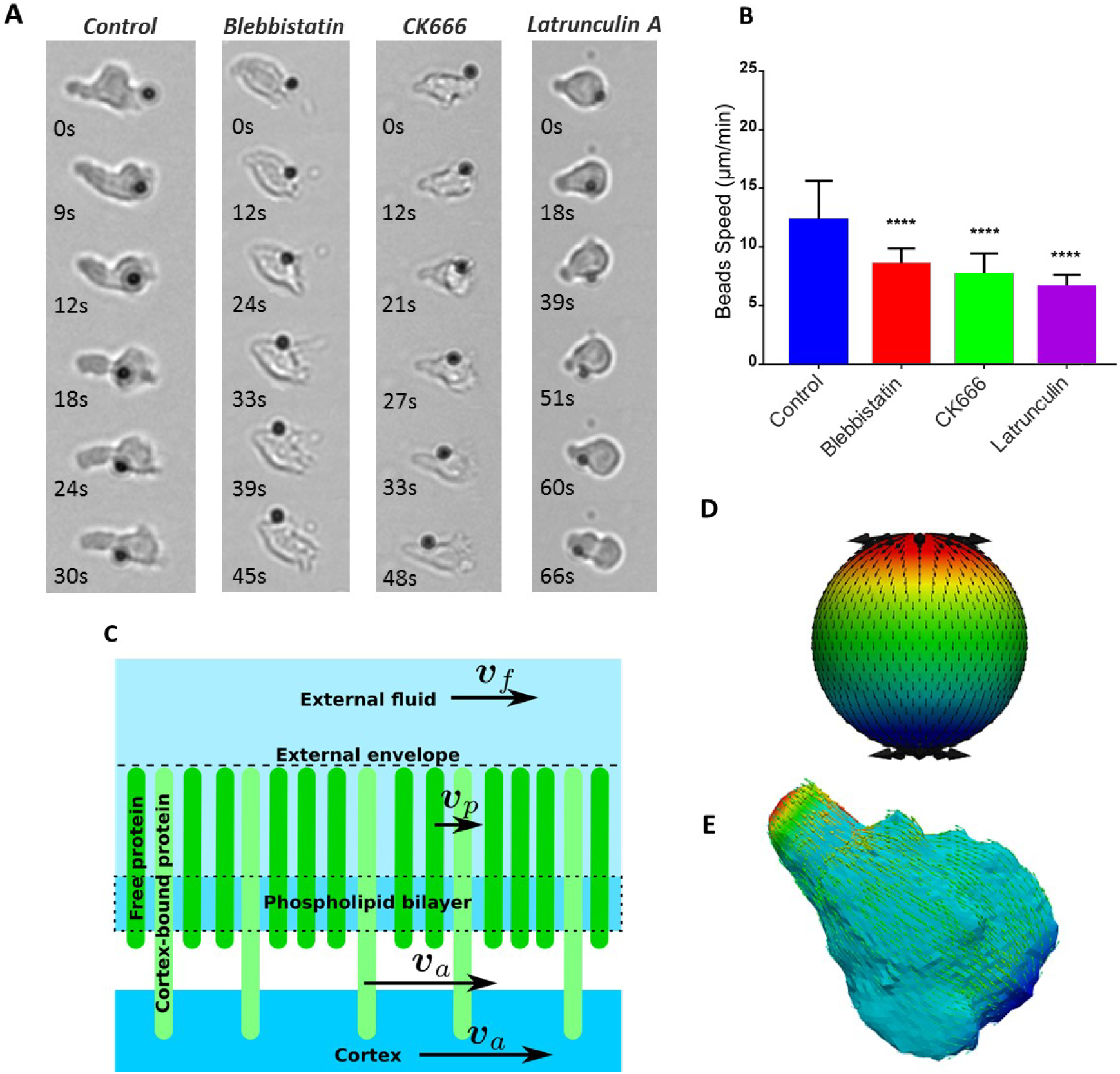
External envelope retrograde flow propels swimming. **(A)** Bright field images of ICAM-coated beads travelling from front to back on the cell membrane of swimming primary human effector T-cell in HBSS control media, and with 50 μM blebbistatin, 100 μM CK666 and 0.05 μM Latrunculin A (left to right). Scale bar 10 μm. **(B)** Travelling speed of ICAM-coated beads versus inhibitors type (n=17 cells for each case, error bar is SD, ****: p<0.0001, Dunnet’s multiple comparison test vs control condition) **(C)** Cartoon of the external and internal structure of the cell envelope considered in the modeling of membrane dynamics. **(D-E)** Retrograde flow pattern on a model spherical cell (D) and a cell with an experimental shape (E) extracted from soSPIM images of (Figure 5-B). Swimming speed is found in both cases equal to the speed of the membrane at equator.

### Retrograde flow of cell envelope can propel swimming

To analyze quantitatively the propulsion strength induced by tangential movements of the cell envelope, we proposed a basic model of retrograde flow for a cell envelope composed of an internal actin cortex, a cytoplasmic lipidic membrane and transmembrane proteins protruding outside the cell (**Figure 5**-C). The transfer of movement from the inner cortex to the fluid surrounding the cell can be total or partial depending on the coupling mechanism between the cortex, the lipidic membrane and the transmembrane proteins, as discussed below. We first considered that the cell surface was made of a homogeneous envelope with an average treadmilling velocity proportional to the velocity of the actin cortex, *v*_*a*_, with a proportionality transmission coefficient, denoted as *β*. In the laboratory frame, the velocity of the fluid adjacent to the cell envelope, denoted as *v*_*f*_, is:

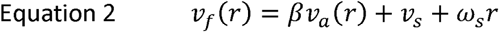

where *v*_*s*_ and *ω*_*s*_ are respectively the translation and rotation swimming velocities of the cell. They were obtained by solving the Stokes equations (see Suppl. Mat.) in the fluid outside the cell, taking as boundary conditions that the flow vanished at large distances away from the cell, and that it obeyed Equation 2 at the cell surface. The obtained flow field was parametrized by the still unknown quantities *v*_*s*_ and *ω*_*s*_, which allowed us to express the viscous forces acting on the cell. The system was closed by imposing that the total force and torque acting on the swimmer vanished, yielding the values of *v*_*s*_ and *ω*_*s*_. A remarkable observation is that the swimming velocity does not depend on the viscosity of the suspending medium for a given cortex velocity (see Suppl. Mat. for a proof), which is consistent with experimental observations (Suppl. Mat. Figure S 1). The problem could be solved analytically for a sphere (**Figure 5**-D) and the swimming velocity was given by *v*_*s*_ *= βv*_0_, where *v*_0_ is the retrograde flow velocity at the equator. For other shapes, we solved numerically the problem using the boundary integral formulation for the Stokes equations (see Suppl. Mat.). In particular, we discretized a shape of T-lymphocyte obtained by 3D spinning disk microscopy (**Figure 5**-E) and introduced an actin source in a small region at the front of the cell and a sink at its rear. The overall flow pattern was similar to the spherical one and the swimming velocity was close to the velocity of the cell envelope in the central region of the cell (in the cell frame), as with the spherical shape (Suppl. Mat. Movie 10). Besides, since most experimental data were obtained for cells close to a rigid wall, we checked numerically that the swimming velocity was largely insensitive to the distance to the wall (Suppl. Mat. Figure S 5). Altogether, this analysis shows that swimming speed has the same magnitude as treadmilling speed of the external envelope, meaning that cortex retrograde flow can propel lymphocytes swimming and may even be the sole propelling source.

### Cell swimming at the molecular scale

The propulsion model by a treadmilling membrane yielded almost equal speed of cell swimming and of envelope treadmilling, but experimental swimming speed, *v*_*s*_ *∼* 5 μm.min^−1^, is more than 2 times smaller than the envelope treadmilling speed, measured above 10 μm.min^−1^ with beads attached to the membrane. This difference suggests an incomplete coupling between the external cell envelope and the surrounding fluid, whereas our model considered a total coupling between the fluid and a homogeneous envelope. In fact, both the composition and the dynamics of a cell cytoplasmic membrane are highly heterogeneous at the molecular level. The external fluid is in contact with the numerous lipids and transmembrane proteins of the membrane, and each of these components have different diffusion coefficients and interactions properties. If the retrograde flow of actin cortex and actin-bound transmembrane proteins is well attested, the circulation of lipids and of non-actin-bound transmembrane proteins is hardly documented. To get more insight into the molecular dynamics of the cell envelope at the time scale of tens of seconds and at the spatial scale of the entire cell, which are the relevant scales for cell swimming, we therefore performed live FRAP-TIRF measurements on non-adherent cells maintained in the vicinity of the probing glass/fluid interface using depletion force induced by addition of dextran in the medium. On RFP-actin transfected cells, we observed the motion of actin clusters that displayed no detectable diffusion. Actin cortical cytoskeleton behaved like a solid gel (**Figure 6**-A,C,E and **Movie** 11 and **Movie** 12) flowing backwards at 24 ± 9 μm.min^−1^ (**Figure 6**-I). This result is consistent with literature data with a speed value in the top range of the ones measured in other cellular systems (between 6 and 20 μm.min^−1^) ^11,22,29,33,34^. For transmembrane proteins, different dynamics are expected whether proteins bind or not to the actin cortex. To shed light on this issue, we used specific fluorescent antibodies to probe an actin-bound protein, the integrin LFA-1 in its high affinity state, and a non-actin-bound protein, the T cell receptor ligand MHC-1. Like actin, actin bound proteins LFA-1 was found to form clusters that persistently flowed backward (**Figure 6**-B,D,E) with an average velocity of 25 ± 5 μm.min^−1^ (**Figure 6**-I). This velocity is similar to the velocity of actin retrograde flow, which is consistent with a strong attachment rate of high-affinity integrins to subcortical actin. No diffusion was detectable. By contrast, non-actin-bound proteins MHC-1 displayed mostly a diffusive dynamics in FRAP experiments (**Figure 6**-G and **Movie** 11) with a characteristic diffusion coefficient of *D =* 0.26 ± 0.22 μm^2^.s^−1^ (**Figure 6**-J). Similarly, for the lipidic layer, FRAP experiments with Vybrant^®^ DiO lypophilic molecules inserted in the cytoplasmic membrane (**Figure 6**-H and **Movie** 11) yielded a diffusion coefficient of 3.1 ± 1.8 μm^2^.s^−1^ (**Figure 6**-J). Altogether, these results clearly revealed that molecular dynamics within plasma membrane differ strongly between the actin-bound and non-actin-bound transmembrane protein, the former being actively dragged backwards and the latter being mainly diffusive. This heterogeneity may explain the discrepancy between the speeds of swimming and actin retrograde flow.

**Figure 6:**
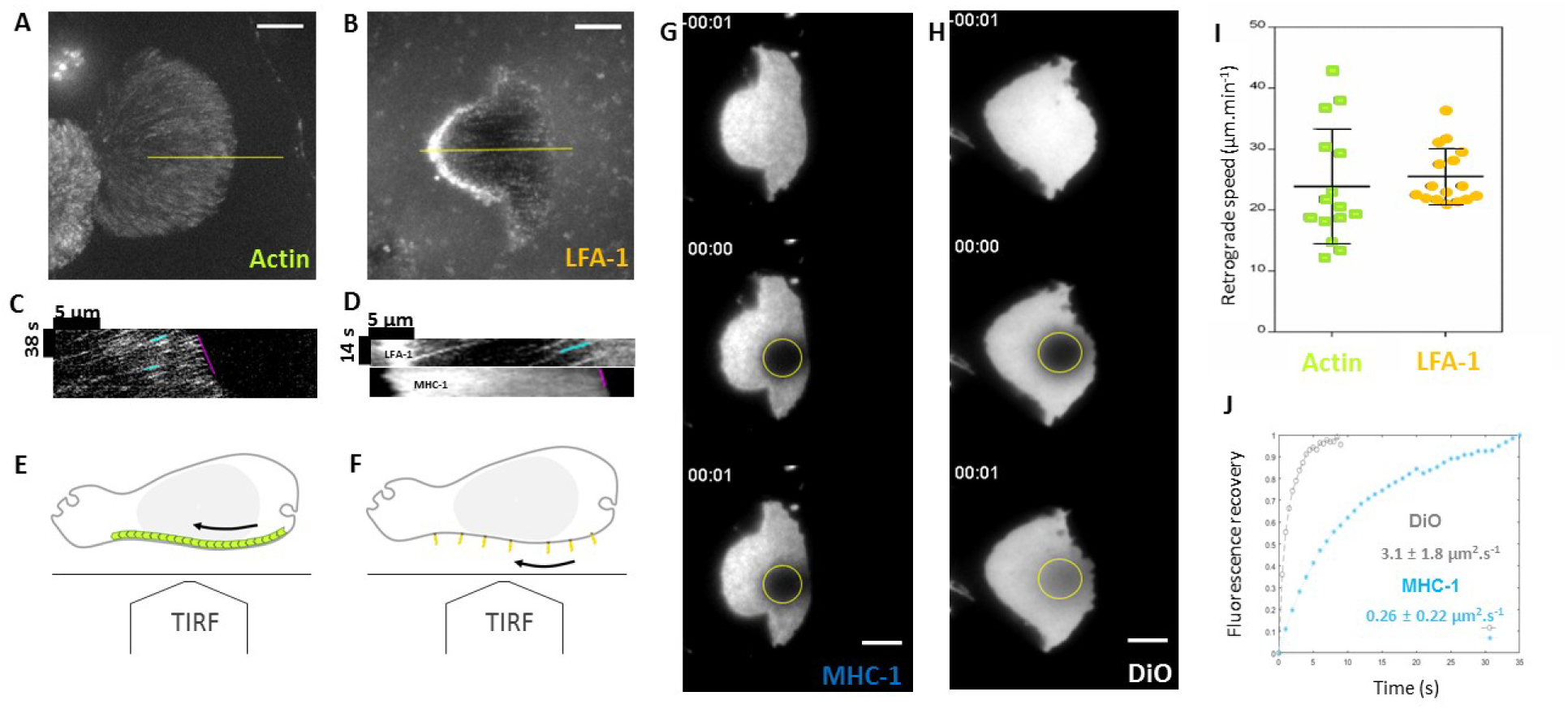
Actin-bound and non-actin-bound transmembrane proteins have different dynamics. **(A,B)** Maximum intensity projection of TIRF sequences on adhesion free cells **(A)** transduced with RFP-actin, **(B)** antibody M24 that binds the actin-bound integrin LFA-1 in its high affinity state. **(C,D)** Representative kymographs along the yellow line in figures A and B with the front advance highlighted in magenta and clusters retrograde flow highlighted in blue. **(E,F)** Schematics representing a side view of a non-adherent cells observed with a TIRF objective and highlighting the retrograde flow of internal actin (E) and external LFA-1 integrins (F). **(G,H)** TIRF-FRAP experiments on adhesion free cells, **(G)** stained with anti-HLA-ABC that binds the non-actin-bound MHC-1 type I proteins, and **(H)** with membrane lipidic marker DiO. Yellow circles are frapped regions used to calculate fluorescence recovery curves shown in (J). **(I)** Speed values for retrograding actin measured by cluster tracking as in (D) (n=7 cells, 51 clusters tracked) and by FRAP as in Suppl. Mat. (n=8 cells) and, and for LFA-1 clusters (n=16 cells, 106 clusters tracked), all normalized to the front of the cell. **(J)** Averaged FRAP curves for DiO (n=10 cells) and MHC-1 (n=14 cells). All values were normalized and corrected by a non-bleached cell. Scale bars 5 μm.

### LFA-1 molecules are extruded at the cell front and MHC-1 are only diffusive

Membrane molecular dynamics is clearly heterogeneous, but the exact participation of the different membrane components to molecular paddling remains unclear. The backwards motion of actin-bound proteins certifies their participation to swimming. In contrast, for lipids and non-actin-bound proteins, their fast diffusion limited the detection of treadmilling speed to values above 50 μm.min^−1^, which backward flow is expected to be smaller than actin speed of 25 μm.min^−1^. To get further insight into the exact traffic properties of non-actin-bound proteins, it is however interesting to consider that a sustainable axisymmetric retrograde flow of material at cell membrane must topologically be coupled to an internal anterograde flow. Mechanism of internal integrins recycling by anterograde vesicular transport have recently been documented in the literature^35–38^, albeit not for leukocytes nor LFA-1. FRAP experiments on cell front revealed indeed a source of unbleached LFA-1 integrins at the cell leading edge (Figure 7-A and Suppl. Mat Movie 13) arising from the extrusion of integrins after internal trafficking. Interestingly, the cycle extrusion/treadmilling of integrins occurs here within a minute, which in consistent with migration and swimming scale, whereas literature reported longer timescales around 15-30 min^36,37^. This observation valid the complete cycling of integrins by treadmilling ad vesicular transport. In contrast, FRAP experiments with MHC-1 proteins displayed a fluorescence recovery only from the cell back (Figure 7-B and Suppl. Mat. Movie 13) without source of material at cell front, which directly supports that internal recycling is not occurring for this non-actin-bound protein. Altogether, FRAP-TIRF data support strongly that actin-bound proteins undergo mostly advected retrograde flow externally and forward vesicular transport internally with an axial symmetry around cell poles, whereas non-actin-bound proteins undergo mostly diffusion at the cell surface without internal recycling. Hence, the cell external envelope does not treadmill as a whole and only a fraction of envelope molecules treadmill and paddle to propel swimming.

**Figure 7:**
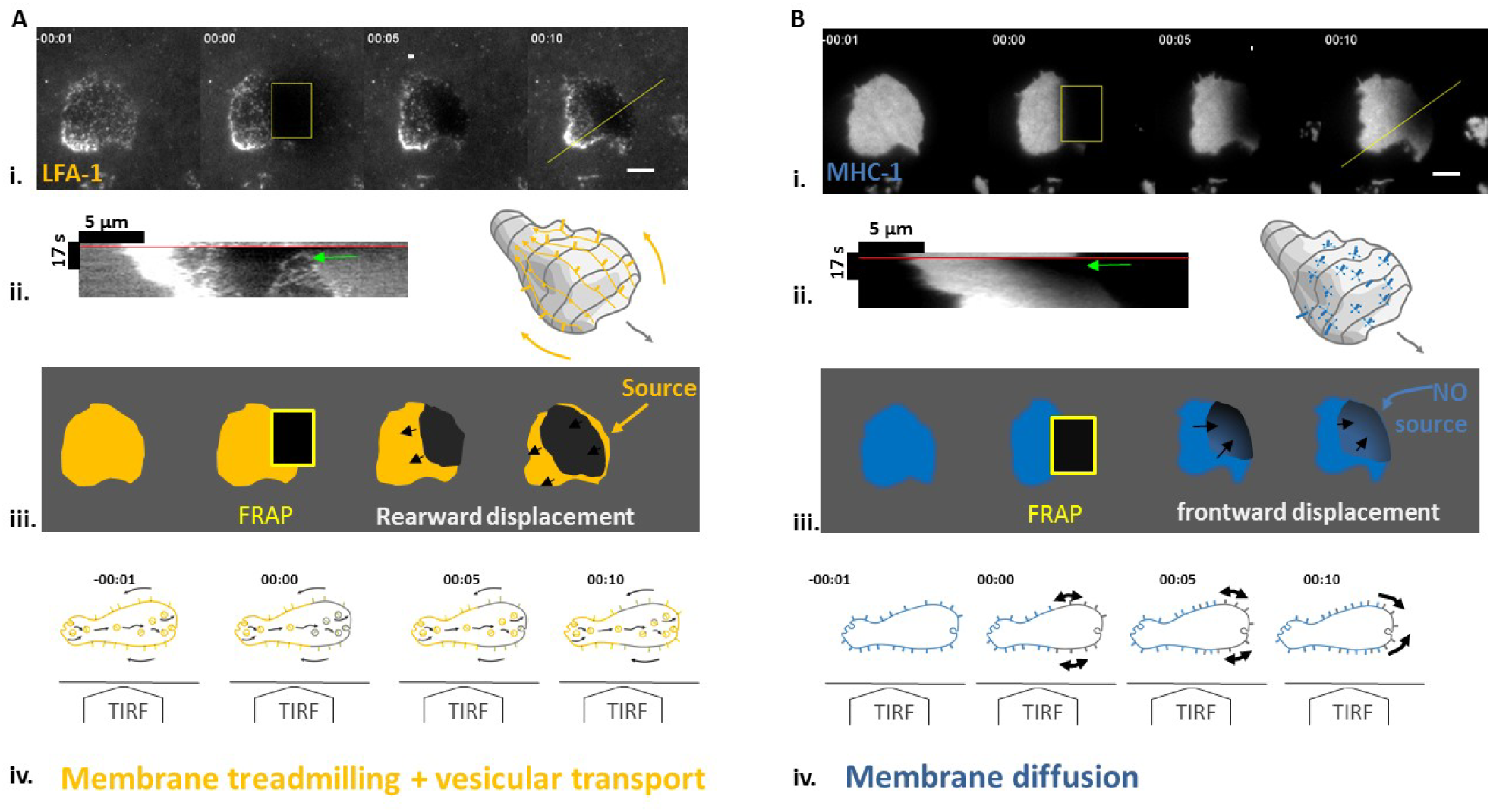
Only actin-bound transmembrane proteins are recycled by internal vesicular transport and external treadmilling advection. **(A)** Staining of high affinity LFA-1 and **(B)** of MHC-1. **(i)** Sequences of TIRF images before FRAP, after frap of cell leading edge, then 5 and 10s after FRAP. Scale bar 5 μm. **(ii)** Kymographs along yellow lines of figures A and B, with red lines indicating FRAP time and green arrows pointing at the cell front shortly after FRAP. 3D cartoon illustrate backward treadmilling of LFA-1 and 2D diffusion of MHC-1. **(iii)** Schematics of experimental results in (i.) illustrating that fluorescence recovers from the cell leading edge for LFA-1, revealing a source at cell front, and from the back for MHC-1 revealing the absence of source at cell front, **(iv)** Side view schematics a non-adherent cells observed with a TIRF objective illustrating the dynamics of transmembrane proteins evidenced in TIRF experiments. For actin-bound LFA-1, source at cell front reveals internal frontward transport of fresh material exocytosed at cell front, whereas for non-actin bound MHC-1, purely diffusive transport dominates (H).

### Model of retrograde flow transmission

In order to model the coupling between actin retrograde flow and the external fluid through a heterogeneous envelope, we considered the layers of transmembrane proteins protruding in the external fluid as a brush of polymeric molecules, and analyzed the flow inside the brush composed of either advected or diffusing molecules. Based on polymer science developments, the brush of proteins was considered as a Brinkman medium^39^ with a hydrodynamic penetration length, denoted as 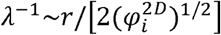, where 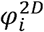 is the area fraction occupied by advected transmembrane proteins (e.g. integrins) and *r* is the typical lateral extent of the proteins on the membrane. The transmission coefficient can be expressed as:

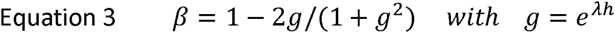

where *h* is the brush thickness. We first considered the effect of advected proteins linked to actin. The two integrins LFA-1 and VLA-4 were measured on T lymphocytes with an average occurrence of 25,000 and 15,000 molecules per cell (Suppl. Mat. Figure S 2). Considering a radius *r* = 5 nm for each integrin and an excess of membrane in microvilli of 150 %, the surface occupancy of LFA-1/VLA-4 corresponds to a mere 0.1 %. The fact that integrins are not in high affinity state at the same time would tend to decrease further this value. However, T lymphocytes express several other integrins than LFA-1/VLA-4 as well as other transmembrane proteins linked to actin (e.g. TCR or CD44), which contribute to increase the fraction of cell surface occupied by advected molecules. All in all, exhaustive experimental data are lacking to determine the exact value of 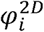. Assuming 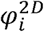 of 2% and *h* of 20 nm, the theoretical coupling value was found at ϐ = 0.2, which is in agreement with the experimental result of a swimming velocity 5 times smaller than the cortex retrograde flow. The effect of diffusing transmembrane proteins was also taken into account by considering that they were indirectly advected by the drag of the external fluid and by viscous interactions within the membrane. Modeling of fluid drag and estimation of membrane viscosity from FRAP measurements allowed us to show that membrane viscosity dominates over the external fluid drag (Suppl. Mat) and that the presence of diffusing transmembrane proteins reduces the value of β (Suppl. Mat Figure S 6).

## DISCUSSION

Amoeboid migration of mammalian cells, i.e. leukocytes and cancer cells, has attracted intensive interest in the last decade for the ubiquitous ability to migrate at high speed in various 2D and 3D environments. The requirement of adhesion with a substrate remains a widely accepted hallmark for 2D migration ^11,20,21,22,5–7,25,40,23^, whereas two studies reported non-adherent motility or swimming with leukocytes^18,41^. In this context, we provided here direct experimental proof and quantification of swimming by wild lymphocytes, together with an original theoretical model of molecular paddling that explains propelling of lymphocytes amoeboid swimming.

In principle, cell swimming without flagella can be propelled either by normal (protrusion) and/or tangential (treadmilling) motion of the cell envelope. Mechanistic studies of swimming by eukaryotic cells have mostly focused on amoeba Dictyostelium discoideum and favored propulsion by shape deformation rather than treadmilling^13,14^. Interestingly, the recent study of O’neil et al^18^ proposed instead that membrane treadmilling propelled the optogenitically-triggered swimming of a macrophages cell line, whereas the contributions of protrusions was not assessed. With wild lymphocytes, we showed experimentally that swimming was barely affected by inhibition of protrusion waves whereas that swimming speed correlated significantly with disruptions of membrane treadmilling. Modeling calculations further proved that protrusions travelling at 10-20 μm/min can not to propel swimming at speeds such elevated as the 5 μm/min observed for lymphocytes. In the end, membrane treadmilling is involved in lymphocytes swimming whereas shape deformations not.

Actin dynamics acts as a hybrid motor of membrane treadmilling, powered by polarization and contractility. Polymerization at the cell front pushes backwards the nascent crosslinked cortical network, while contractility at the cell rear pulls backwards the network. It is not clear if the two mechanisms combine their action on a continuous actin gel or if they act independently on two disconnected actin networks at the cell front and rear. However, we find here that swimming relies more on frontal actin polymerization than on rear myosin-II contractility. This observation diverges from recent reports on adherent crawling ^21,40,42^–^45^ and swimming ^18^ cells that attribute a dominant role to gradient of contractility across cells. Our results are in turn consistent with studies proposing that contractility is marginally involved in propulsion and mostly relevant for detachment of cell rear (for Dictyostellium amoebae)^46^ and squeezing of cell nucleus through pores (for dendritic cells)^5^. In the end, it remains that membrane treadmilling is propellnig leukocytes swimming, whereas membrane treadmilling itself may be powered either by actin network contractility^18^ or actin polymerization depending on cell type.

Hydrodynamic coupling between a treadmilling membrane and a surrounding fluid is the key of amoeboid swimming, but its characteristics at molecular level have not been precisely considered experimentally nor theoretically. Studies on motility usually consider that cellular membranes treadmill as a whole, which theoretically yields a ratio between swimming and membrane speeds ranging from 2/3 to l^47^. O’neil et al ^18^ supported the hypothesis of membrane treadmilling as whole is supported because they measured swimming speeds equal to 2/3 of membrane speeds and they observed front-rear gradients of several membrane components (lipids, and proteins). It can however be argued that correlations between membranes and cells speeds relied on a limited data set and that gradients of proteins are not a proof of a direct molecular treadmilling. In contrast, we find on a large cohort of cells, and after extraction of diffusive motion, that swimming speed of lymphocyte is significantly too low to arise from propulsion by a homogeneous membrane treadmilling; swimming speed was 1/3 of membrane treadmilling speed (measured with a bead attached to the membrane) and 1/5 of the actin-bound proteins speed. Furthermore, a direct measurement of transmembrane proteins dynamics revealed a strong heterogeneity in membrane treadmilling; some molecules treadmilled backwards at high speed and were recycled internally towards cell front, whereas others were diffusive and not recycled internally toward cell front (Figure 8). Finally, theoretical modelling supports quantitatively that heterogeneous treadmilling of a cell membrane can account for the slowness of cell swimming as compared to actin treadmilling.

**Figure 8:**
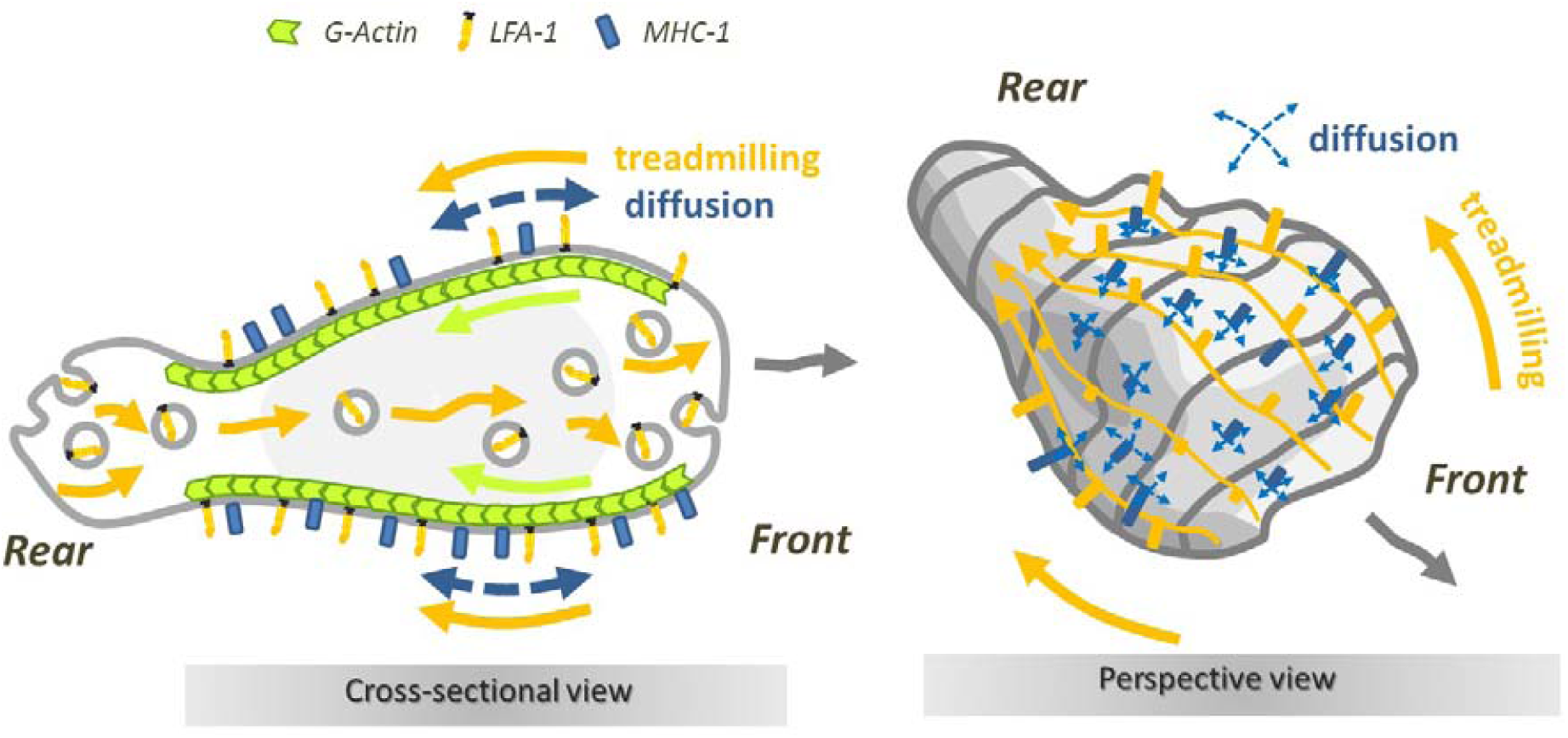
Slow swimming is modeled by a partial coupling of fast actin treadmilling motion to a fraction of transmembrane proteins recycled from rear to front by internal transport. **(A)** Cross sectional view. The complete cycling of actin-bound proteins (here LFA-1, yellow) comprises endocytosis at cell rear, internal forward advection by vesicles, exocytosis at cell front and advection at cell membrane by linkage to retrograde actin flow (green). Non-actin bound proteins (here MHC-1, blue) are mostly diffusing at cell surface. (B) Perspective view of a swimming cell with actin-bound proteins advected backwards (yellow), entangled with non-actin bound proteins. Molecular paddling resulting from interactions between external fluid, and actin-bound and non-actin-bound proteins yields a speed significantly lower than actin according to our theoretical model and experimental data.

Our swimming model sheds also new light on the ubiquitous motility of amoeboid cells. The adaptation of mammalian amoeboid cells to various environments has repeatedly been attributed to their capability to switch between different migration modes versus the environment^9,20,22,28,28,40,48^. The mode of adherent crawling is generally attributed to a sequence of cell front protrusion/attachment of and cell rear pulling /detachment, whereas the more intriguing mode of non-adherent migration in 3D has been explained by various mechanisms depending on cell types^8,22^: blebbing in the cell front and transfer of actin cytoskeleton into the novel bleb^43,49,50^, intercalation of protrusion into gaps and discontinuities of the matrix to advance like on a ladder^51,52^, chimneying via active gel pushing-off the wall^10^, cell rear contractility stabilizing a single bleb conformation^21^, water permeation throughout the cell body^53^ or treadmilling coupled to friction with the substrate^8,11^. In our case, there was no evidence of a mode switching between crawling and swimming sequences on patterned substrates. But the adaptation of lymphocytes to various microenvironments may also not require several modes^9^. The single action of envelope treadmilling can indeed mediate ubiquitous migration of T lymphocytes in a 3D matrix, on a 2D substrate and in suspension in a fluid, with only slight difference detectable in terms of speed. As experimentally measured, an adhesive environment provides strong cell/substratum coupling and cell speeds close to actin treadmilling, whereas a liquid environment provides a partial cell/liquid coupling (due to membrane heterogeneity and not to environment fluidity) and speeds significantly lower than actin treadmilling.

In the end, actin retrograde flow can drive ubiquitous amoeboid migration within fluid and solid, and with or without non-adherent, however, the physiological role of swimming remains enigmatic for leukocytes. Since the traffic of immune cells (or invasivity of cancer cells) relies mostly on matrix-associated migration and swimming is only relevant for planktonic eukaryotic cells (like amoeba), it is probable that the basic machinery for swimming is an evolutionary conserved ability that has found novel functions for ubiquitous crawling of mammal cells.

## Supplementary information

### Supplementary movies

Movie 1: **From Crawling to Swimming in the vicinity of a substrate.** (Left) Crawling on adhering ICAM-1-treated substrate. (Right) Swimming on Pluronic^©^ F127 treated surface. First sequence, 20x bright field transmission microscopy, then 63x bright field transmission microscopy and finally x63 reflected interference contrast microscopy.

Movie 2: **3D imaging of primary human effector T cell swimming in spinning disk microscopy.** Videomicroscopy sequence of swimming T cells stained with CMFDA (5-chloromethylfluorescein diacetate) on Pluronic^©^ F127 treated surface for 14 min 40 sec with a time lapse every 20 s and 10 slices every 1 μm. Some unpolarized cells do not swim. The arrow points a polarized and swimming cell that crosses the whole field of view. Scale bars in μm indicated on axis, magnification: 63X.

Movie 3: **Immediate transition between crawling and swimming.** Migration on alternative 40 μm wide stripes of adherent ICAM-1 and non-adherent Pluronic^©^ F127 prepared by LIMAP^31^. Superposition of fluorescent image (ICAM-1, red), bright filed transmission image (greyscale) and reflection interference contrast microscopy image (bright green corresponds to cells adhesion fingerprints) taken at 63x. Scale bar 20 μm.

Movie 4: **Swimming in bulk suspension.** Movie of two cells suspended in a medium of matched density using a microscope tilted by 90°and a flow cell oriented vertically for sideway observations.

Movie 5: **Role of actomyosin network in swimming motion.** Movies in bright field at 63x of primary human effector T cells swimming on an antiadhesive substrate in the presence of actin inhibitors: control, 50 μM blebbistatin, 100 μM CK666 or 0.05 μM Latrunculin. Scale bar 20 μm.

Movie 6: **3D imaging of primary human effector T cell swimming in so-SPIM mode.** Video microscopy movie of RFP-Lifeact transfected cells, showing lamellar-protrusion forming with random orientation in cell front and travelling backwards at around 10 μm.min”^−1^.

Movie 7: **Rearward travelling shape deformations.** Bright field videomicroscopy at 63x of primary human effector T cells swimming on an antiadhesive substrate and showing protrusion retrograde motion, nucleus squeezed forward through constricted rings. Scale bars 10 μm.

Movie 8: **Swimming of a cell by the motion of two blebs on the cell surface.** The color here represents the mean curvature of the cell surface.

Movie 9: **Cell envelope retrograde flow revealed by attached beads.** Bright field videomicroscopy of ICAM-coated beads travelling from front to back on the cell membrane of swimming T-cells in HBSS control media, 50 μM blebbistatin100 μM CK666 or 0.05 μM Latrunculin A. Scale bars 50 μm.

Movie 10: **Numerical simulation of swimming by retrograde flow.** Cell shape is extracted from experiments. The swimming is shown in the laboratory frame. Color code on the surface represents the production/consumption of the cortex material. Small spheres are fictitious tracers moving with the cortex velocity. Transmission coefficient *β =* 1.

Movie 11: **Molecular analysis of the dynamics of cell external envelope by TIRF-FRAP.** TIRF-FRAP experiments on primary human effector T-cell (Top left) transfected with a GFP-Actin by lentiviral infection, (Top right) stained with membrane lipidic marker DiO, (Bottom left) stained with antibody Mab24 that binds an actin-bound protein, the integrin LFA-1 in its high affinity state, and (Bottomright) stained with anti-HLA-ABC that binds the non-actin-bound MHC-1 type I proteins. Scale bars 5 μm.

Movie 12: **Cytoskeleton retrograde flow.** TIRF imaging of a primary human effector T lymphocyte transfected with GFP-Actin and displaying backward travelling of clusters. Scale bar 5 μm.

Movie 13: **Evidence of internal recycling at the cell front for advected protein LFA-1 and not for diffusive protein MHC-1.** TIRF-FRAP experiments on primary human effector T-cell (Right) stained with antibody Mab24 that binds the actin-bound proteins LFA-1 in high affinity state, and (Left) stained with anti-HLA-ABC that binds the non-actin-bound proteins MHC-1. The cell front is frapped and fluorescence recovers only from the front for LFA-1, in agreement with an internal vesicular recycling of integrins from back to front, and only from the rear for HLA, in accord with a surface diffusion mechanism.

## Supplementary information on experimental results

### Experimental swimming velocities are independent of medium viscosity increase up to 100 times

Hydrodynamic interactions between an amoeboid lymphocyte and a fluid are sufficient to promote momentum transfer. In order to test if the viscosity of the medium influences the efficiency of cell-fluid coupling, we performed swimming experiments in culture medium supplemented with dextran of molecular weights 2,000 kDa to increase its viscosity up to 100 times. Viscosity of solutions were measured on a Bohlin Gemini 150 rheometer equipped with cone-plate geometry (cone angle 20°, diameter 60 mm) at T=22°C. The change of osmotic pressure and change of viscosity (Figure S 1-A) had no significant effect on swimming velocity in the explored range (Figure S 1-B). This observation is consistent with predictions of the model for swimming propelled by cell envelope retrograde flow.

**Figure S 1:**
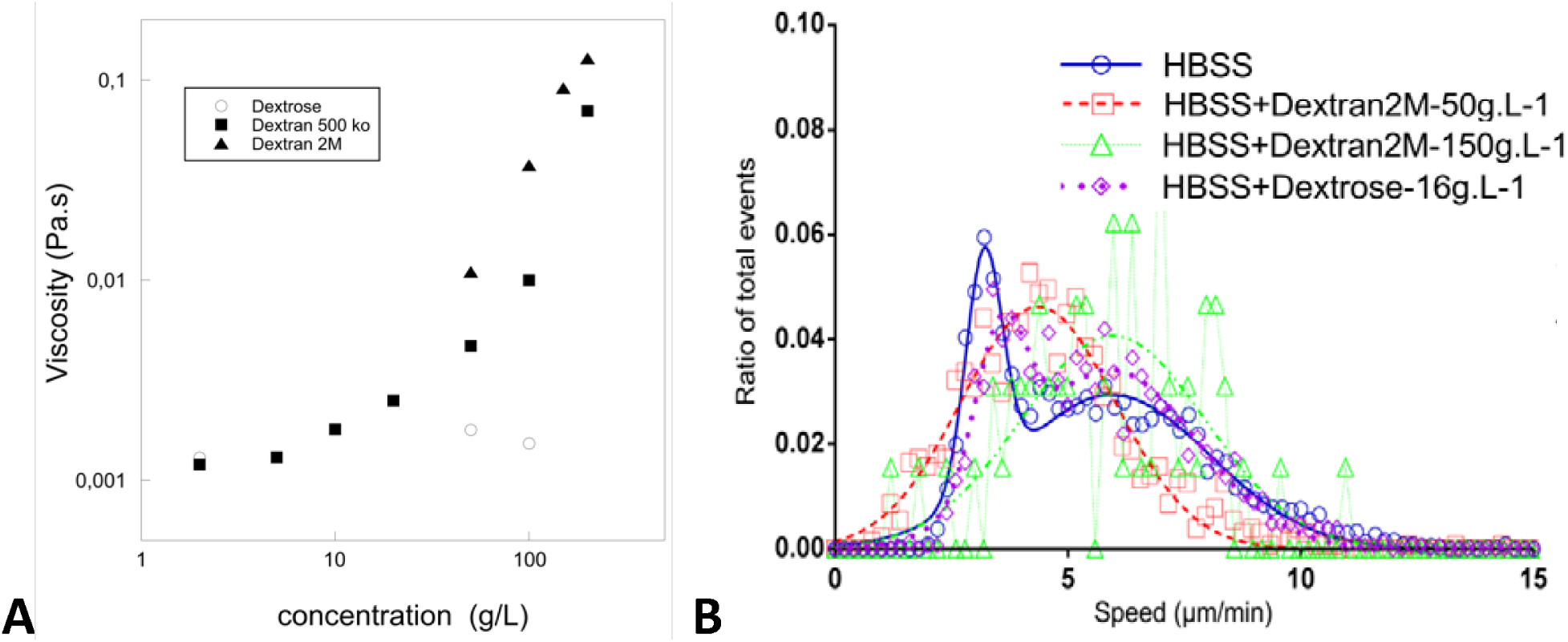
Cell speed is independent of the medium viscosity. **(A)** Viscosity of Dextran and dextrose solutions in HBSS versus concentration. **(B)** Histogram of raw curvilinear velocities for cells in normal medium and medium with viscosity increased 10 x (Dextran 2,000 kDa at 50g/L) and 100 x (Dextran 2,000 kDa at 150g/L) and osmotic control condition (Dextrose 16g/L). Data for increased viscosity cases are fitted with a single average speed of 4.4 μ m min ^−1^ and 6.0 μ m min^−1^ for 10x and 100x increased viscosity condition respectively. Data for osmotic control are fitted with a double Gaussian with an average speed of 5.9 μ m.min ^−1^ for the active swimming cells. Ncells = 4342 (HBSS), 1262 (Dextran 50 g.L^−1^), 64 (Dextran 150 g.L ^−1^), 1449 (Dextrose); Nexperiments = 5 (HBSS), 3 (Dextran 50 g.L^−1^), 3 (Dextran 150 g.L^−1^), 2 (Dextrose).

### Quantification of LFA-1 and VLA-4 expression on effector T cells

Quantification LFA-1 and VLA-4 number per cell was performed by quantitative cytometry (Figure S 2) and yielded an average number per cell of 25000 for LFA-1 and 13000 for VLA-4.

**Figure S 2:**
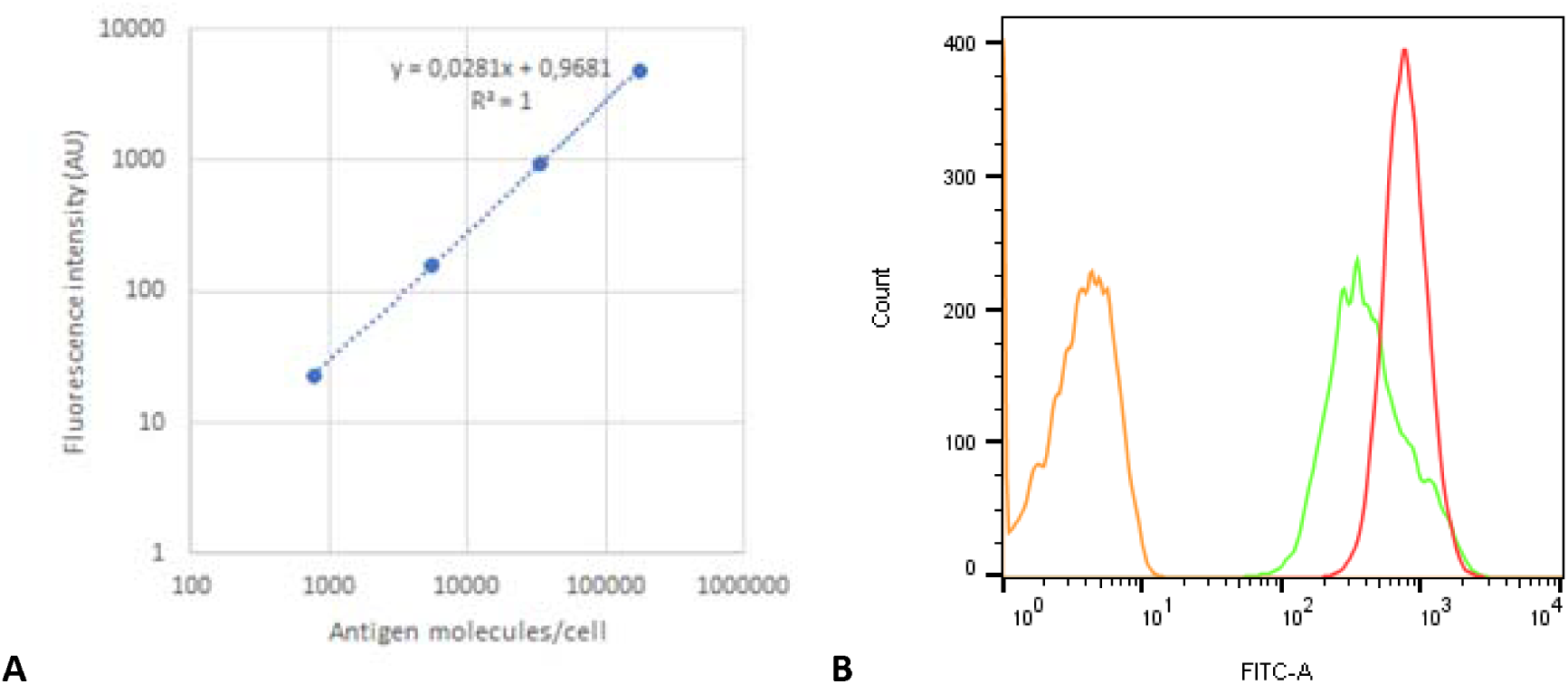
(A) Calibration curves with the secondary antibody and calibration beads (CellQuant calibrator kit, ref 7208, Biocytex) (B) fluorescence histograms of T cells stained by indirect immunofluorescence with specific monoclonal antibodies (CD49d (HP2/1) for VLA-4 and CD11a (Hi111) for LFA-1).

## Supplementary Table

## Supplemental information on theory and simulations

### Cell diffusion and persistent swimming: Model

We model the motion of the cells as a combination of deterministic swimming and random noise. The swimming velocity *v*_*s*_ *= v*_*s*_*p* is assumed to have a constant absolute value but the orientation vector p can vary in time. The random noise here consists of translational diffusion with diffusion coefficient *D*_*t*_ and rotational diffusion with angular diffusion coefficient *D*_*r*_. This noise accounts both for thermal fluctuations and for the active dynamics of the cell. Since cells are swimming close to a wall, the dynamics of orientation p and position r of the swimmer are effectively two-dimensional. The model described above belongs to a broad class of persistent random walk problems, which have enjoyed a lot of attention in the literature^55^. We therefore give here only a brief overview of the solution process and the final expression of the mean square displacement of the cell as a function of time and the model parameters.

### Cell diffusion and persistent swimming: Solution

We first calculate the correlation 〈*p*(*t*_0_) · *p*(*t*)〉. The probability density function Ψ(*p,t*) for the swimmer to have orientation *p* at time *t* satisfies the following equation:

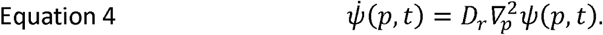

where *∇*_*p*_ = (*I* – *p* ⊗ *p*) · *∂*_*p*_ is the gradient operator on a unit circle representing possible orientations of p, and *I* is the identity matrix. The right hand side of Equation 4 expresses the angular diffusion process. If *p*(*t*_0_) = *p*_0_ the initial condition for Equation 4 is

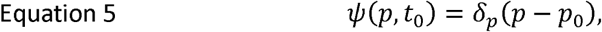

where *δ*_*p*_ is the Dirac function. Equation 4 is solved by expanding *ψ*(*p,t*) in Fourier harmonics of *p*, which represents the eigenfunctions of the Laplace operator 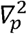:

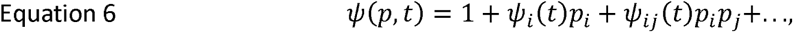

where *ψ*_*ij*_(*t*) and so on are symmetric and traceless. The quantity of interest here is *ψ*_*i*_ = 2〈*p*_*i*_ (*t*)〉. Substituting Equation 6 into Equation 4 yields *ψ*_*i*_(*t*)= *ψ* _*i*_ (*t*_*0*_)*exp*[*D*_*r*_(*t*_*0*_ – *t*)] for *t* ≥ *t*_0_, resulting in

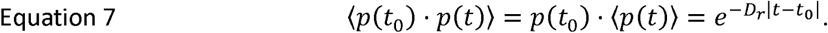

The displacement due to persistent motion is calculated by integrating Equation 7:

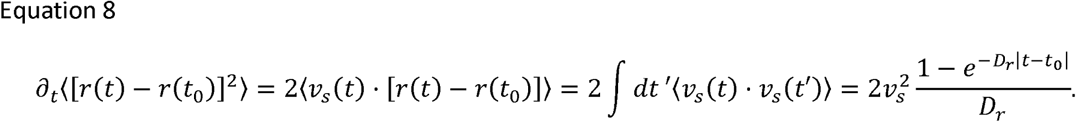

Integrating Equation 8 and adding the contribution of the translational diffusion yields

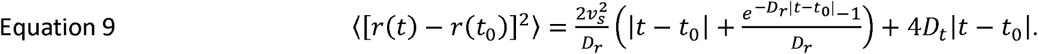

Equation 9 reduces to Equation 1 for *tD*_*r*_ ≪ 1.

### Model of swimming by wave of shape deformation

In order to simulate the bleb-driven swimming of the cells, we model the cell as an elastic capsule, to which active forces are applied, while maintaining the zero net force and torque conditions (Figure S 3). Application of the active force density concentrated in small regions of the cell surface results in the formation of bleb-like protrusions. Further, the location at which the active forces are applied is moved with a prescribed velocity *v*_*bleb*_ along the surface of the cell. In addition to the magnitude of the active force and the bleb velocity, the life-time of the blebs *T*_*bleb*_, which describes the duration for which the active force is applied, is also an important parameter. This application of time dependent active force leads to formation and movement of the blebs along the cell surface. The swimming velocity is obtained by solving the hydrodynamic problem in fluids inside and outside the cell, as described below.

**Figure S 3:**
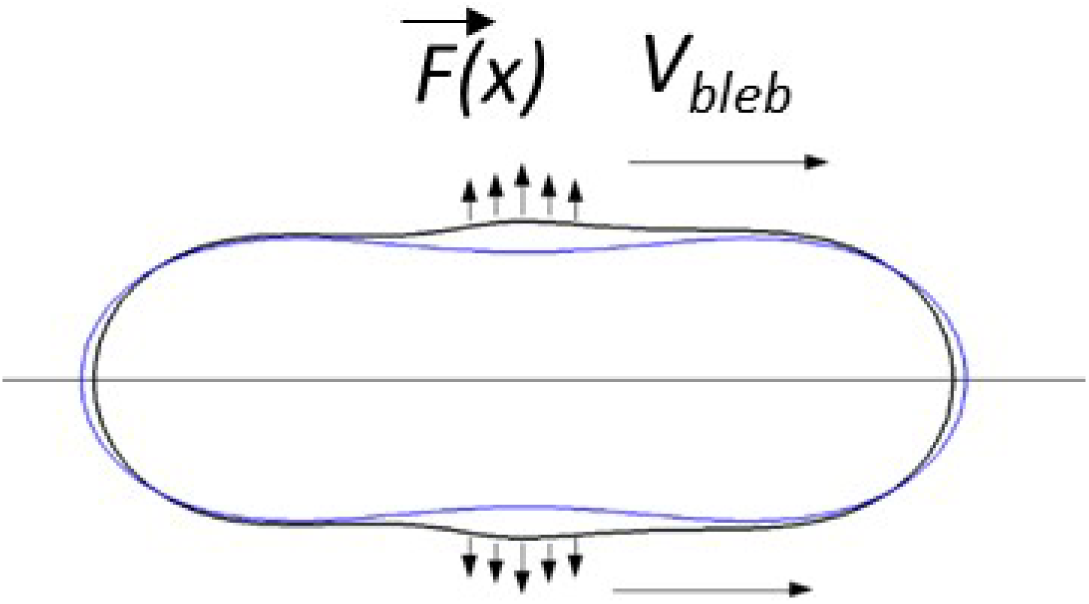
Schematic of a swimming cell by protrusive blebs. Blue and black contours are the initial and deformed configurations of the cell, respectively.

The configuration of the cell during the course of a bleb formation and motion is shown in Figure 4-D. The maximum size of the blebs for different values of active force amplitudes is shown in Figure S 4 and and the velocity of the cell by the bleb motion in Figure S 4-B. For all the simulations *v*_*bleb*_ was fixed at 0.02 and force amplitude *F* was varied. The plot shows that in bleb-driven swimming, the cell velocity *v*_*cell*_ 10^−3^*v*_*bleb*_. Furthermore, the dependence of the swimming velocity on the force amplitude is *vF*^2^.

**Figure S 4:**
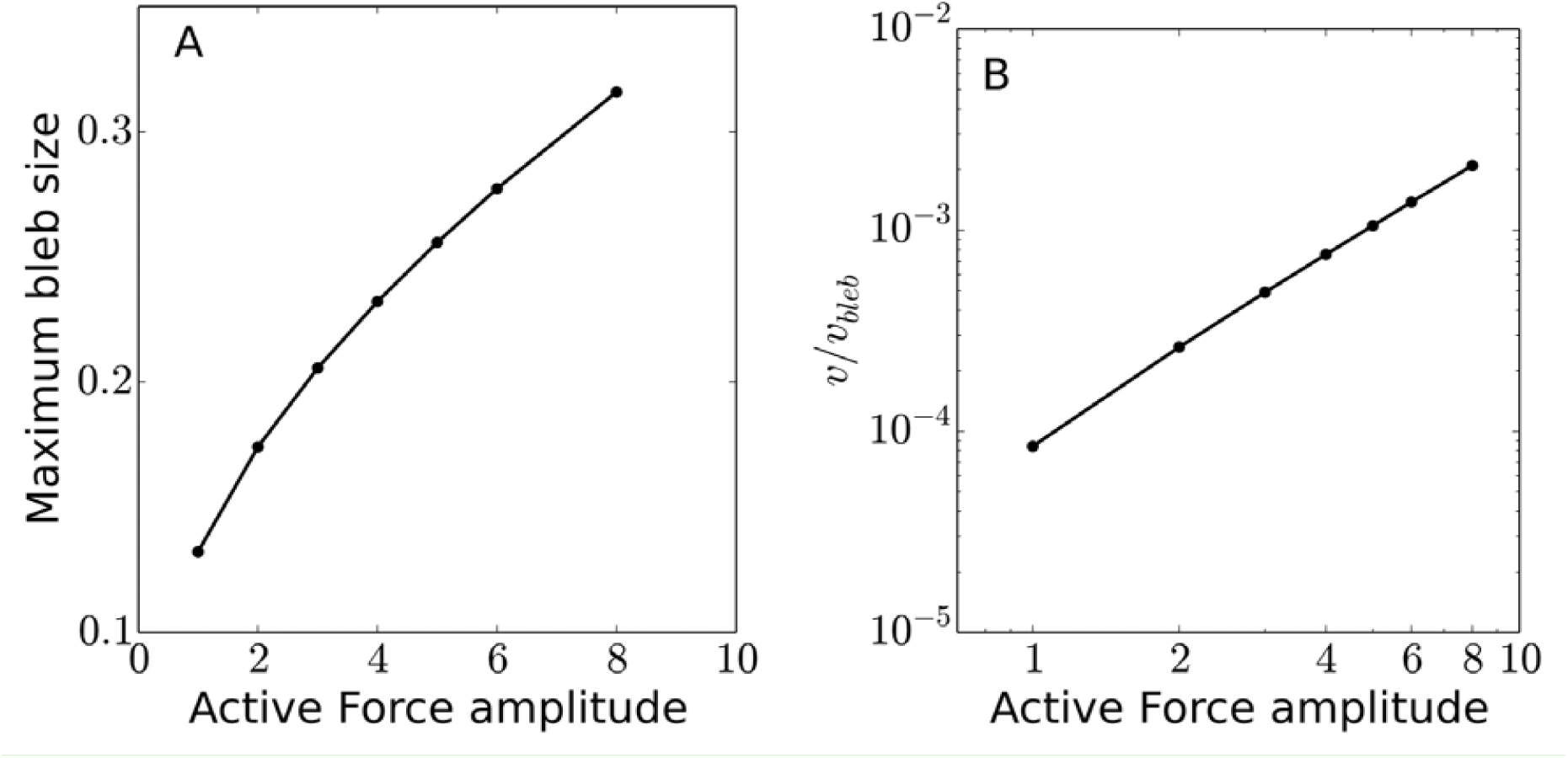
(A) Maximum size of the blebs for different values of active force amplitudes (see Figure S 3). (B) Dependence of average cell velocity *v*_*cell*_10^−3^*v*_*bleb*_ on the active force amplitude. Here the bleb velocity relative to the cell center has been kept fixed at *v*_*bleb*_ *=* 0.02.

### Model of swimming by envelope retrograde flow: Actin cortex flow

Actin polymerization expands the cortex in the front region of the cell, whereas myosin-induced contractility and actin depolymerization consume the cortex in the rear region of the cell. Between these regions, the actin moves from front to back along the cell membrane. An overpressure of the cortex at cell front due to actin accumulation and an underpressure at cell rear due to myosin-induced contraction can both contribute to forces driving the retrograde flow of the cortex. We thus model the driving force as the pressure gradient along the cell surface, ∇^*S*^*P*, where *P* is the pressure field and *∇*^*s*^ is the surface gradient operator. This gradient implies a net cortex flow field *v*_*a*_(*r*) which, in a simple approximation, can be taken locally proportional to the driving force:

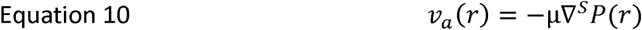

Where *μ* is a mobility coefficient. Here we assume that the shape is fixed so that the flow is tangential to the surface in the reference frame comoving with the cell. By choosing an appropriate pressure field, Equation 10 can be proven to be exact for any axisymmetric flow (see below). It is interesting to note that Equation 10 is identical to the Darcy law, valid in a porous medium or a Hele-Shaw flow. The cortex region could be viewed as a thin layer, a curved Hele-Shaw geometry, hence Equation 10 can be inferred from classical hydrodynamics. Equation 10 is closed by the mass conservation equation:

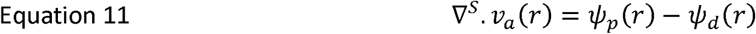

Here we express the change of the local 2D concentration of actin (per cortex area) due to the advection by the flow (the left hand side) through the local rate polymerization *ψ*_*p*_(*r*) (source) and the local rate of depolymerization *ψ*_*d*_(*r*) (sink), *ψ*_*p*_(*r*) and *ψ*_d_(*r*) are zero everywhere except in a small domain localized in the front and rear regions of the cell, respectively. Substituting Equation 10 into Equation 11 gives an equation for *P* which can be solved. Once *P* is determined the actin flow field *v*_*a*_(*r*) can be obtained from Equation 10.

### Model of swimming by envelope retrograde flow: actin flow in an axisymmetric case

The purpose is to show that for axisymmetric cells Equation 10 holds automatically. The cell shape and the actin velocity can be parametrized in cylindrical coordinates as

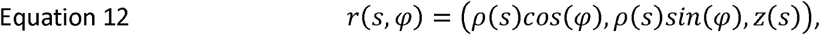

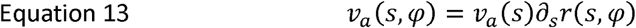

where *s* is the arclength measured from the front pole of the cell, *φ* is the polar angle, *ρ*(*s*) and *z*(*s*) are shape functions, and *∂*_*s*_*r*(*s, φ*) is the tangent vector along the meridian. The properties of *s* are such that |∂_*s*_*r*(*s*, *φ*) | = 1. Equation 10 can be easily verified if we define *P*(*s*, *φ*) as

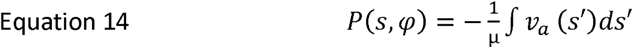

## Model of swimming by envelope retrograde flow: Exact solution for a spherical swimmer

The actin flow can be obtained explicitly for a spherical cell having a point source of actin *ψ*_*p*_(*r*) *=* 2*Rv*_0_*δ*(*r* – *r*_*N*_) at the North Pole *r*_*N*_ and a point sink ψ_*d*_(*r*) *= 2Rv*_0_*δ*(*r* – *r*_*s*_) at the South Pole *r*_*s*_. Here *R* is the sphere radius and *v*_*0*_ is a constant having a dimension of velocity that turns out to be equal to the retrograde flow velocity at the equator in the cell frame. We present the solution in spherical coordinates. Defining the origin as the center of the swimmer and the polar direction as source point of actin, we write the velocity at a point (*Rsinθcosφ, Rsinθsinφ, Rcosθ*) as

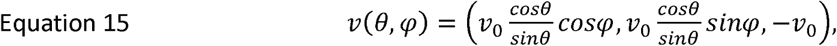

where *θ* is the azimuth angle. The corresponding actin pressure is written as

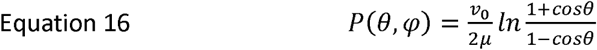

The swimming velocity is then given by *v*_*s*_ *= βv* _0_

### Flow outside the cell

Since the Reynolds number in the problem is extremely low, the flow *V*_*f*_ in the fluid outside the cell envelope satisfies the Stokes equations:

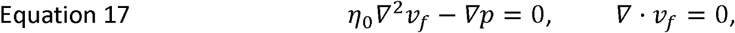

where *η*_0_ is the viscosity of the suspending medium. Equation 17 are solved together with the boundary conditions given by *V*_*f*_ = 0 at infinity and Equation 2 of the main text. The unknowns *v*_*s*_ and *ω*_*s*_ are solved for from conditions:

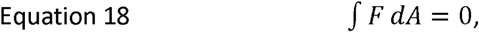

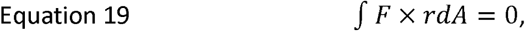

where *F* is the surface force density applied locally by the fluid to the cell, *dA* is the area element and integrals are taken over the boundary of the cell envelope. Equation 18 and Equation 19 express that no external forces or torques act on the swimmer. The forces *F* can be expressed through the viscous stress tensor σ of the fluids

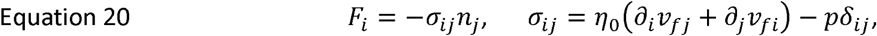

where *n* is the outward normal to the boundary of the cell. It follows from the linearity of Equation 17 and Equation 2 of the main text that for any *a*, if *v*_*f*_, *v*_*s*_, *ω*_*s*_, *σ, F*, and *p* represent a solution of the problem for a given cell shape, *v*_*a*_ and external fluid viscosity *η*_0_, then *v*_*f*_, *v*_*s*_, *ω*_*s*_, *aσ, aF*, and *ap* represent a solution of the problem for the same shape, *v*_*a*_ and external fluid viscosity *aη*_0_. This implies that changing viscosity of the suspending medium does not affect the swimming velocity for the same velocity of the retrograde flow and the same transmission coefficient *β*. This result is consistent with experimental observations (cf. Suppl. Mat., Figure S 1 and “Experimental swimming velocity are independent of medium viscosity increase up to 100 times”).

### Model of swimming by envelope retrograde flow: Numerical method for any swimmer

We parametrized the surface of the cell obtained in experiments by a triangular mesh. The Laplace equation for *P* (Equation 10 substituted in Equation 11) was solved by finding a stationary solution of a diffusion equation. The flow in the suspending fluid was solved for using the boundary integral formulation. The details of the numerical procedure and the validation are given in ^56^.

### Model of swimming by envelope retrograde flow: vicinity of a wall is negligible

We have also solved the numerical problem for a swimmer near a solid wall. The no-slip boundary condition at the wall was imposed by taking a modified Green’s function in the boundary integral formulation, as discussed in Pozrikidis C (1992)^57^. The same shape and retrograde flow field were taken as in the unconfined case. The orientation of the wall was chosen consistently with the experiment but the position was varied in order to scan different gaps between the cell and the wall. The resulting dependence of the swimming velocity (assuming *β* = 1) is shown in Supplemental Figure S5.

**Figure S 5:**
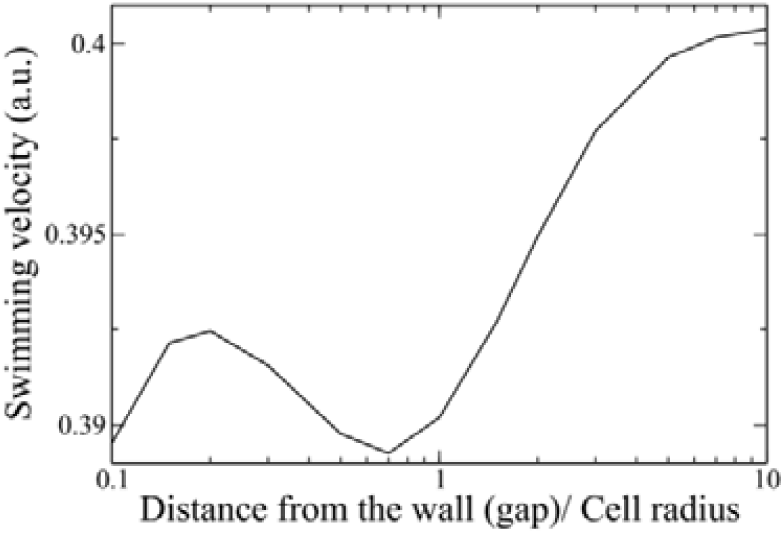
Calculated swimming velocity of cell powered by cell envelope treadmilling versus the distance of the cell to the wall normalised by the cell size. The presence of the wall influences only marginally cell speed between 0.39 and 0.4.

### *Model of swimming by envelope retrograde flow:* Molecular paddling model

The purpose of this section is to introduce a detailed model of the transfer of the cortex retrograde flow to the fluid surrounding the cell. The model is based on a mean-field approximation: we consider a region of a cell boundary that, on the one hand, is large compared to the size of individual proteins, and, on the other hand, is sufficiently small compared to the cell scale. These assumptions allow us to consider the cell boundary to be flat and to represent all relevant quantities as a function of the distance from the cortex, averaging them over the two remaining coordinates. The freely diffusing and the cortex-bound proteins are thus modeled as a homogeneous porous medium with an effective viscous friction with the fluid. A similar model was considered^58^ for a flow of fluid inside a brush of polymers covering a wall and subject to an external flow. We therefore only list here the main ingredients of the solution and final results.

The following analysis is written in the reference frame comoving with the phospholipid bilayer. We consider the actin-bound proteins to move with the cortex velocity *v*_*a*_, as suggested by our measurements showing that high affinity integrins LFA-1 are advected at a speed very close to the one of actin cortex (Figure 6-I), and we call v_p_ the speed of proteins not bound to actin. The exact numbers of advected and non advected proteins is however not known precisely. Finally, we assume that the individual protein molecules interact only via hydrodynamic fields in the outside fluid, thus excluding short-range solid friction between them. The proteins also interact via hydrodynamic fields inside the bilayer. However, since the net velocity of the bilayer is zero in the chosen frame, this interaction would only result in a correlation of velocities in pairs of proteins located closely to each other. This would represent a higher-order effect in the concentration of cortex-bound proteins than the one considered here.

The protein brush can be modeled by the Brinkman equations^39^, which can easily be motivated as follows. The fluid in the brush is to leading order function only of the coordinate z orthogonal to the cell membrane (since the brush thickness is small as compared to the cell size), and obeys the one-dimensional nonhomogeneous Stokes equation

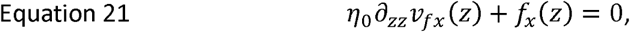

Where *f*(*z*) is the volume-related force density applied by the brush on the fluid, given by

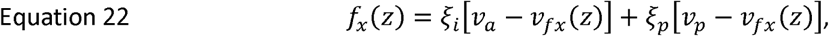

where 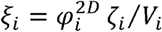 is the volume-averaged drag coefficient of actin-bound proteins (e.g. integrins LFA-1 in the experimental part) and 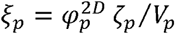 is the volume-averaged drag coefficient of passively advected proteins (e.g. MHC-1 in the experimental part). Here *φ*^2*D*^ is the area fraction of corresponding proteins, *ζ* is the corresponding viscous drag coefficient, and *V* is the volume of the extramembrane part of the corresponding protein.

The velocity of the free proteins, *v*_*p*_, is calculated by requiring the sum of drag forces applied by them on the fluid and the bilayer to be equal to zero, which gives a condition:

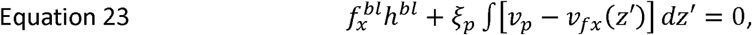

where *h* is the thickness of the brush and *h*_*bl*_ is the thickness of the bilayer. The drag experienced by the freely advected protiens from the phospholipids of the bilayer 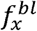 is expressed as

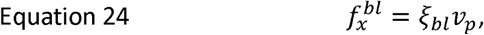

where 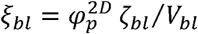 the volume-averaged drag coefficient inside the bilayer for passively advected proteins, *ζ*_*bl*_ is the corresponding Stokes drag coefficient for one protein, and *V*_*bl*_ is the volume of the protein part inside the bilayer. Note that the cortex-bound proteins experience drag inside the bilayer just as the passive ones do but their velocity is fully determined by the flow of the cortex, to which they are firmly attached.

Equation 21 and Equation 22 can be solved for *v*_*fX*_ as a function of *v*_*a*_ and *v*_*p*_ by using the two boundary conditions:

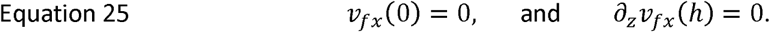

Equation 23 and Equation 24 yield the expression of *v*_*p*_ as a function of *v*_*a*_, and thus the full expression of the fluid velocity field as a function of drag coefficients, the membrane thickness and *v*_*a*_. From this knowledge we determine *β* (which is the transmission coefficient of the cortex flow to the fluid at the brush surface, *z = h*). *β* is function of the drag coefficients and the viscosities. The explicit forms of *β, v*_*p*_, and *V*_*fx*_ read

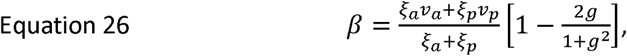

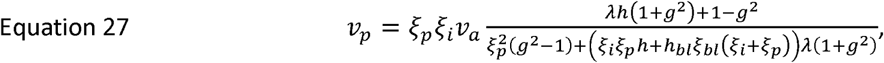

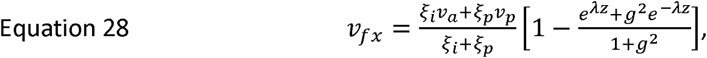

where *λ*^2^ = (*ξ*_*i*_+*ξ*_*p*_)*/η* _0_and *g = e*^*λh*^.

The Stokes drag coefficients of the proteins in the outer fluid are written as ζ = *6nR*_*h*_*η*_*0*_ *≈* 10∼^−10^ *kg/s*, where *R*_*h*_ is the Stokes radius, which we take here as 6 nm for simplicity. The Stokes drag coefficient in bilayer can be estimated directly from the measurements of the diffusion coefficient 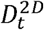 of MHC-1 freely advected proteins : 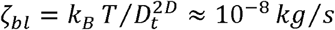. The thickness of the bilayer is taken as 8 nm. The brush thickness is taken as 20 nm, which we estimate from the length of integrins in activated state. The volume of the external part of the proteins is taken as 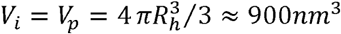. The volume of the bilayer segment of the passively advected proteins is estimated as 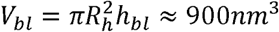. Supplemental Figure S 6 shows the transmission coefficient *β* and *β*_*p*_ *= v*_*p*_*/v*_*a*_ as a function of the concentration 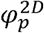.

**Figure S 6:**
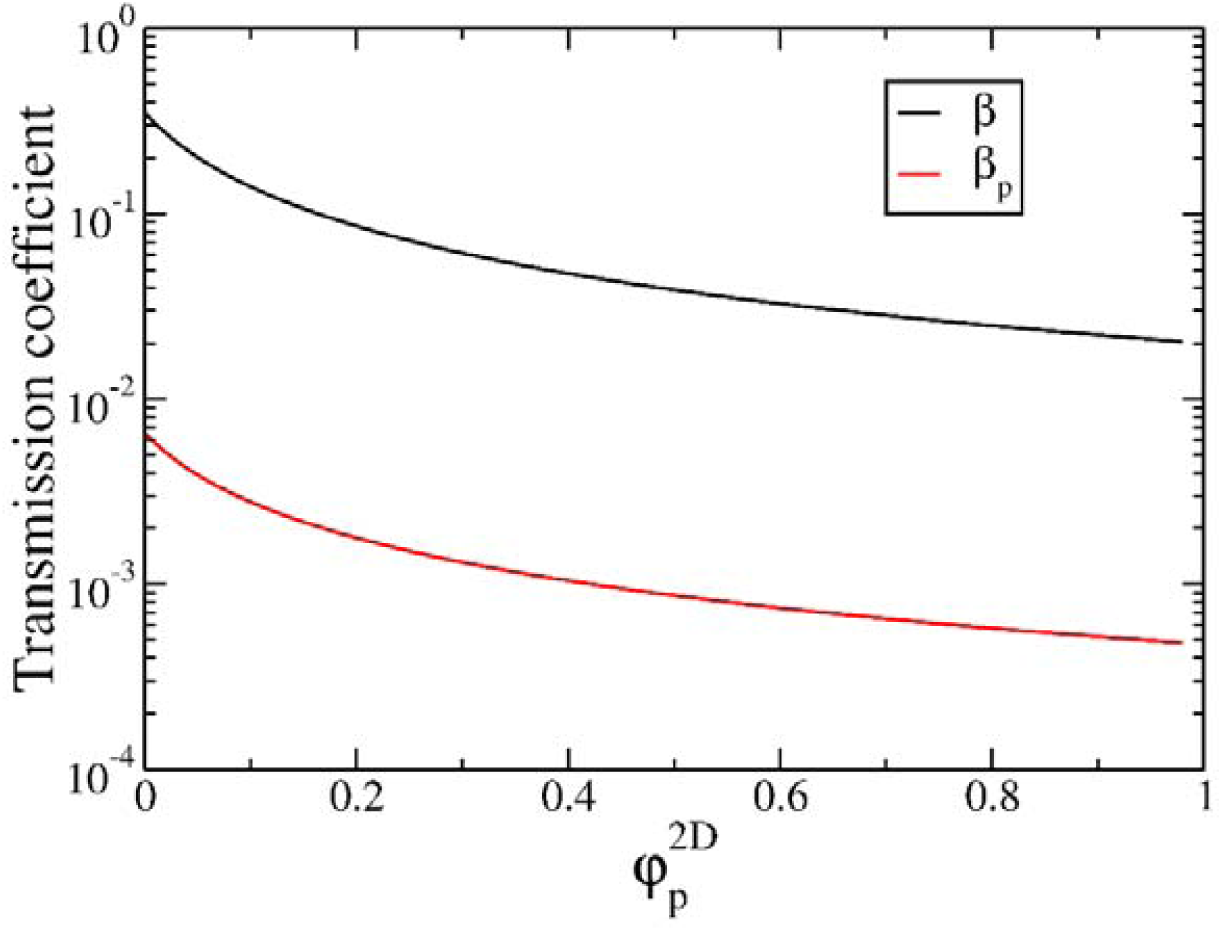
Transmission coefficients of the cortex flow to the fluid *β* (black curve) and to non advected proteins *β*_*p*_ (red curve) as a function of the concentration 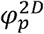, as given by Equation 26 and Equation 27. *R*_*h*_ = 6*nm, η*_0_ *=* 0.001*Pa*·*s, h =* 20*nm*, 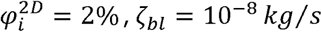.

The analysis above was performed in the reference frame of the phospholipid bilayer. The experimental results do not allow us to establish whether a significant retrograde flow of the phospholipids is present in the reference frame of the cell envelope. Assuming the average local velocity of the phospholipids *v*_*bl*_ in the reference frame of the cell envelope is known, we can express the velocity fields in the reference frame of the swimmer envelope as *V*_*f*_ *+ v*_*bl*_ for fluid velocity, as *v*_*p*_ *+ v*_*bl*_ for freely diffusing proteins and as *v*_*a*_ *+ v*_*bl*_. The full expression for transmission coefficient is then written as 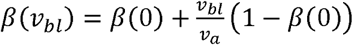, where *β* (0) is given by Equation 26 and *v*_*a*_ is measured in the reference frame of the cell envelope. This shows that allowing for retrograde flow of the bilayer further increases the transmission coefficient.

## Material and methods

### Cells

Whole blood from healthy adult donors was obtained from the “Établissement Français du Sang”. Peripheral blood mononuclear cells (PBMCs) were recovered from the interface of a Ficoll gradient / “Milieu de separation des lymphocytes” (eurobio, les Ulis, France). T cells were isolated from PBMCs with Pan T cell isolation Kit (Miltenyi Biotec, Bergisch Gladbach, Germany), then were stimulated for 48⍰h with anti-CD3/anti-CD28 Dynabeads (Gibco by Thermo Fischer Scientific, Waltham, MA) according to the manufacturer’s instructions. T lymphocytes were subsequently cultivated in Roswell Park Memorial Institute Medium (RPMI) 1640 (Gibco by Thermo Fischer Scientific, Waltham, MA) supplemented with 25⍰mM GlutaMax (Gibco by Thermo Fischer Scientific, Waltham, MA)), 10% fetal bovine serum (FBS; Gibco by Thermo Fischer Scientific, Waltham, MA) at 37°C, 5% CO2 in the presence of IL-2 (50⍰ng/ml; Miltenyi Biotec, Bergisch Gladbach, Germany) and used 6 to 10⍰days after stimulation. At the time of use, the cells were >99% positive for pan-T lymphocyte marker CD3 and assessed for activation and proliferation with CD25, CD45RO, CD45RA and CD69 markers as judged by flow cytometry.

### Quantitative cytometry for integrin expression level

For the quantification, we used the CellQuant calibrator kit (ref 7208, Biocytex). T cells were stained by indirect immunofluorescence with specific monoclonal antibodies, CD49d (HP2/1) for VLA-4 and CDlla (Hilll) for LFA-1, then analyzed by quantitative flow cytometry. The expression level of the tested antigen was determined using the kit calibration beads.

### Transduction of cells

For soSPIM experiments with LifeAct transduced cells, virus was produced in HEK 293T cells by co-transfecting the lentiviral plasmids pLenti.PGK.LifeAct-Ruby.W (a gift from Rusty Lansford - Addgene plasmid #51009) with psPAX2 and pMD2.G (a gift from Didier Trono - Addgene plasmid #12260 and #12259). PBMC were transduced by spinoculation of virus using polybrene, after 48h activation with CD3-CD28 Dynabeads. The cells were then cultured with IL-2, and used 8 days after activation. The expression of LifeAct-RFP was controlled by flow cytometry. For TIRF-FRAP experiments cells, RFP-Lentivirus for RFP-actin transduction were bought from Merck (Lentibrite™ RFP*-β*-actin lentiviral biosensor) and cells were transduced 48h after activation with an MOI of 10. For GFP-actin transfection, plasmid EGFP-Actin-7 from Addgene (ref 56421) was used with the electroporation program Amaxa T20.

### Microfluidic channels and surface treatments

PDMS microchannels were fabricated using standard soft lithography. A positive mould was created with a negative photoresist SU-8 3000 (Microchem) on silicon wafers (Siltronix), then replicas were moulded in polydimethylsiloxane (PDMS) elastomer (Sylgard 184, Dow Corning) and sealed on glass cover slides via plasma activation (Harricks Plasma). The device is composed of one channel with one inlet and one outlet punched with a 2.4 mm puncher (Harris Uni-Core). For adherent crawling experiments, Ibidi channels IV^0.4^ (Clinisciences) were coated overnight at 4°C with 10μg/ml human ICAM-l-Fc (R&D Systems) in Phosphate Buffer Solution (PBS) (Gibco). Channels were subsequently blocked with a solution containing 2.5% bovine serum album (BSA) (w/v; Axday, France) and 2.5% Pluronic acid F-108 (w/v; BASF, Germany) in PBS for 30 min at room temperature, then rinsed three times with PBS and finally with HBSS. Cells were injected at densities around 1.5×10^6^/ml and allowed to equilibrate for 10 min at 37°C before image acquisition. For non-adherent migration or swimming experiments, Ibidi channels IV^0.4^ and PDMS microchannels were incubated with Pluronic F-127 (Sigma-Aldrich) for 30 min at room temperature, then rinsed three times with PBS and finally with HBSS. Cells were injected at a densities around 0.75×10^6^/ml in Ibidi channels and 6×10^6^/ml in PDMS microchannels of height 40 μm. Cells were allowed to equilibrate for 10 min at 37°C before image acquisition.

### Cell treatments

Stock solutions of blebbistatin (Fischer Bioblock Scientific), CK666 (Sigma-Aldrich) and Latrunculin (L12370, 2.37 mM; Molecular Probes) were prepared in DMSO following manufacturer’s specification, stored at −20°C and then diluted in culture medium for used in experiments. Cells were resuspended in solutions of 50 μM blebbistatin, 100 μM CK666 and 50 nM Latrunculin, injected in the microchannels, and allowed to settle in the channels for a period of 30 min at 37 °C before image acquisition.

### Viscosity and osmolarity measurements

Viscosity changes were performed using Dextran of average molecular weight of 1500-2800 KDa (Sigma-Aldrich) at concentrations of 50 and 150 g/L. HBSS alone has a viscosity value of 0.001 Pa.s while the viscosity for HBSS+50g/L Dextran is 0.01 and for HBSS+150g/L Dextran it is 0.1 Pa.s. Adding Dextran to the media increased the viscosity as well as the osmolarity up to 355 mosm/kg for the solution HBSS+150g/L Dextran. Dextrose (Sigma-Aldrich) was then used as an osmolarity control in HBSS media supplemented with 25mM HEPES. Osmolarity measurements for the different media were performed using the automatic Micro-Osmometer Type 15 (Loser Messtechnik), calibration was done using standard solutions of 300 and 900 mosm/kg H_2_O according to the manufacturer’s instructions.

### Experimental fluidic setup

All experiments were performed in a home-made chamber precisely thermostated at 37°C to limit temperature instability potentially inducing flow drifts within fluidic devices. For swimming close to a surface, we used Ibidi channels for experiments in HBSS, Dextrose and 50g/L Dextran solutions, and 40 μm high PDMS microchannels for experiments in in 150g/L Dextran 40 μm high to limit the observation range in the z axis because cells did not sediment. To minimize flow, channels were sealed with the plastic cap for Ibidi channels, or with a 250 μM thickness PDMS film for the PDMS microchannels and the devices on the microscope stage were surrounded by a 100% humidity chamber to minimize evaporation through PDMS. For experiments of swimming in suspension, cells were resuspended in 66% Ficoll to limit sedimentation effects, and injected in 100 μm high channels. Minimization of drifts for swimming in suspension was more challenging than for the swimming close to a substrate. The microfluidic channel was set vertical (along the gravity axis) and the whole microscope was tilted by 90° to get side-observation view. The channel was connected to a microfluidic flow control system (Fluigent MFCS-EZ) to control the unidirectional flow towards the bottom, and we used 2 meter long tubes of 0.5 μm internal diameter to further limit drift by hydraulic resistance. Cell motion were recorder for at least 100 frames every 10 seconds.

### Cell motion imaging

Experiments were performed with an inverted Zeiss Z1 automated microscope (Carl Zeiss, Germany) equipped with a CoolSnap HQ CCD camera (Photometrics) and piloted by μManager ^1.4^. Different objectives were used for bright-field mode (Plan-Apochromat 20x/0.8, 63x/1.4 objectives) and for reflection interference contrast microscopy (RICM) mode (Neofluar 63/1.25 antiflex). A narrow band-pass filter (λ=546 nm ± 12 nm) was used for RICM. Three dimensional imaging was performed on cells stained with a lipophilic tracer DiO (Invitrogen) and cells transfected with lifeAct-RFP cells. The imaging was done using a Spinning disk (Inverted Nikon Eclilpse TI) equipped with two cameras (Photometrics EMCCD evolve) and controlled by Metamorp, and a home made single-objective selective plane illumination microscopy (soSPIM) set-up (see special section for details).

### soSPIM imaging and analysis

The soSPIM system, for single-objective Selective Plane Illumination Microscope, is a recently developed architecture which enables to combine the advantages of low photo-toxicity and high optical sectioning of light-sheet microscopy techniques with the high sensitivity provided by high numerical aperture objectives^59^. The set-up is composed of a high numerical aperture objective (CFI Plan Apochromat VC 60x WI 1.27NA), a beam steering unit and dedicated micro-fabricated devices containing mirrors angled at 45° alongside micro-wells. The soSPIM components are mounted on a conventional inverted microscope (Nikon Ti-E). The micro-fabricated chambers (see ^59^ for detailed descriptions of the chambers) are placed on an axial translation piezo stage (Mad City Lab) within a controlled environment chambers (Tokai Hit) for live cell imaging. Fluorescence emission is collected through the same objective used for excitation and is captured on a sCMOS camera (ORCA-Flash 4.0 V2, Hamamatsu). The whole acquisition process is steered under MetaMorph environment (Molecular Device) using a home-made designed plugin which synchronize the excitation and acquisition processes. Further details of soSPIM setup, calibration, and synchronization are described in^59^. The 3-dimensional time series data sets acquired with the soSPIM set-up were analysed using the freely available software UCSF Chimera^60^ (developed by the Resource for Biocomputing, Visualization, and Informatics at the University of California, San Francisco (supported by NIGMS P41-GM103311)). This software enables to render surfaces of equal fluorescence intensity as well as normalization and alignment of whole 3D time series which enhances the possibility to visualize cell membrane movement in our case.

### Molecular motility imaging

For TIRF-FRAP experiments, cells were resuspended in HBSS-Dextran 150g/L solution at a concentration of 4.5 10^6^ cells/ml, in the presence of CDlla/CD18 (Biolegend, clone M24) and HLA-A,B,C (Biolegend, clone W6/32) primarily conjugated antibodies. Alternatively cells were labeled with Vybrant^®^ DiO by 10 min incubation at 37°C in the presence of 5 ul of dye per 1.5 10^6^ cells, then washed twice with HBSS and resuspended in HBSS-Dextran. For LifeAct-RFP cells, no further staining was required. Cell suspensions were loaded into the devices and centrifuged for 3 min at 200 RCF. Cells were allowed to equilibrate for at least 10 min at 37°C before image acquisition. Movies were recorded on a Nikon Eclipse Ti microscope, equipped with iLas2 system and controlled by Metamorph software. For DiO and MHC-1 staining, diffusion coefficients were calculated using the SimFRAP ImageJ plugin*. LFA-1 cluster speed values were calculated from kymographs performed along the cell axis, while Actin flow was calculated measuring the displacement of the frapped region. All values were corrected by the advance of the front edge, measured by a kymograph along the cell axis, to obtain a value relative to the cell front.

### Cell tracking

For swimming experiment in the vicinity of a substrate (in 2D), cells were tracked with a home-made program (MATLAB software, The MathWorks, Natick, MA, USA) and raw curvilinear velocities of swimming cells were calculated using trajectory time points every 30s. Residual flow drift was corrected on each cell trajectory using the mean x- and y-movements values of all cells between 2 pictures. For high viscosity experiments, the fraction of cells squeezed toward the substrate by depletion force were discarded from the analysis. For swimming experiment in suspension (in 3D), a stack of bright-field images was taken every 10 s across the 100 μm height of the channel with a spatial pitch of 5 μm. To determine the position a particular cell on the x-axis at a given time, we analyzed the intensity distribution of the image of this cell on all images of the x-axis stacks. The best focus corresponded to the image with the minimum standard deviation of the intensity, which yielded a x-position with a precision of 2.5 μm. Each cell trajectory was fragmented in 30s steps and cell-step speed was calculated using coordinates along x- and z-axis. The speed component along y was considered negligible because we selected cells with an orientation perpendicular to Y-axis. Total cell speed was calculated as the mean of all the 30s steps-speed for each cell.

### Beads advection experiments

Streptavidin-coupled beads with a diameter of 2.7 μm (Dynabeads^®^ M-270 Streptavidin, Invitrogen) were washed three times with 0.1% BSA (w/v), then incubated with 0.5μg/ml biotin-coupled Protein-A (Sigma-Aldrich) for 1 hour under stirring at room temperature and rinsed with 0.1% BSA. The beads were then incubated with 500μg/ml ICAM for 2 hours at room temperature and rinsed with 0.1% BSA. A final concentration of 0.125mg/ml Dynabeads was added to the cell suspension. Bright-field images (Plan-Apochromat x20/0.8) were taken every 3s. Beads were tracked manually from the moment the bead attached to the cell front until it reached the cell rear. Cell are moving in the frame of the laboratory and cell rear was taken as a reference of bead position. All experiments were performed at least in triplicate for each substrate and/or drug.

## Contributions

LA and PN worked on all experiments and analysis. AF and CM developed all modeling and simulations, MSR performed modeling of swimming by shape deformation, NGS performed experiments of beads tracking and all FRAP-TIRF assays; XL performed migration assays on patterned substrates, SD and CH prepared cells transfected with GFP actin, MB cultured cells performed transfection with RFP-Lifeact and quantitative cytometry, MPV participated to experiments and analysis, RG and JBS performed 3D live imaging by soSPIM, CH and SD prepared GFP-actin cells, SR performed viscosity measurements and participated to discussions and, OT participated to all experiments and analysis, CM and OT designed the study and wrote the paper.

## Acknowledgments

The work at LAI was supported by the ANR grant RECRUTE, LABEX INFORM, the Région PACA, Institute CENTURI, Nanolane company and Alveole company. We thank the France Bioimaging Platform, funded by the French Agence Nationale de la Recherche (ANR–10–INBS–04–01, ≪investments for the futures ≫). We are grateful to Alphée Michelot for his support and advices with TIRF-FRAP imaging, to Laurence Borge for assistance with the use of the Cell Culture Platform facility (Luminy TPR2-INSERM), to Claire Chardes and Pierre-Francois Lenne for help with light sheet imaging, Michael Sixt for advices on Actin flow measurements, and Yannick Hammon for helpful discussions. Authors also thank the Electronic imaging centre of Bordeaux Imaging Centre for metallization of the soSPIM chambers, the National Research Agency (ANR) grant soSPIM, and the National infrastructure France-BioImaging supported by the French National Research Agency (ANR-10-INBS-04). The work at LIPHY was supported by CNES (Centre National d’Etudes Spatiales), ESA (European Space Agency) and the French-German university programme “Living Fluids” (grant CFDA-Q1-14). C.M. and S.R. thank D.K. Dysthe for many valuable discussions and for having clarified several issues of the amoeboid swimming problem during his one year stay in Liphy in 2011, as well as Alain Duperray and Nawal Quennouz for valuable discussions regarding their preliminary experiments on swimming of neutrophils.

